# Polyamines regulate cell fate by altering the accessibility of histone tails

**DOI:** 10.1101/2024.07.02.600738

**Authors:** Maya Emmons-Bell, Abby Sundquist, Megan R. Rabon, Grace Forsyth, Sylvie Oldeman, Shine Ayyapan, Angeliki Gardikioti, Roshni de Souza, Jonathan Coene, Maryam H. Kamel, Harrison A. Fuchs, Varun S. Sudunagunta, Edna M. Stewart, Steven Verhelst, Joanna Smeeton, Aaron D. Viny, Catherine A. Musselman, Juan-Manuel Schvartzman

**Affiliations:** Columbia University, Herbert Irving Comprehensive Cancer Center, New York, NY, USA; Yale School of Medicine, Boyer Center for Molecular Medicine, New Haven, CT; Department of Genetics and Development, Columbia University Irving Medical Center, Columbia University, New York, NY, USA; Weill Cornell Medical College, New York, NY, USA; Barnard College, New York, NY, USA; Department of Biochemistry and Molecular Genetics, University of Colorado Anschutz Medical Campus, Aurora, CO, USA; University of Leuven, Cellular and Molecular Medicine, Leuven, Belgium; Columbia Stem Cell Initiative; Department of Rehabilitation and Regenerative Medicine; Columbia University Vagelos College of Physicians and Surgeons; Department of Biochemical Engineering, Columbia University; Columbia University Irving Medical Center, Department of Medicine, Division of Hematology/Oncology, New York, NY, USA

## Abstract

Polyamines are polycationic alkyl-amines abundant in proliferating stem and cancer cells. How these metabolites influence numerous cellular processes remains unclear. Here we show that polyamine levels decrease during differentiation and that inhibiting polyamine synthesis leads to a differentiated-like cell state. Polyamines are enriched in the nucleus, where their loss drives changes in chromatin accessibility and histone post-translational modifications. Polyamines interact electrostatically with DNA on the nucleosome core, freeing histone tails to conformations accessible to chromatin-modifying enzymes. Consistent with their role in increasing histone-tail accessibility, polyamines are able to replace MYC’s role in reprogramming to pluripotency. These data reveal a mechanism by which an abundant metabolite influences chromatin structure and function in a direct but sequence independent manner, facilitating chromatin remodeling during reprogramming and limiting it during fate commitment.

## Introduction

The polyamines putrescine, spermidine and spermine are polycationic (positively-charged) alkyl-amines present at high concentrations in eukaryotic cells^1,2^. Polyamines are synthesized by decarboxylation of the amino acid ornithine by the enzyme ornithine decarboxylase (ODC1) (Fig. 1A). At approximately 1 mM, polyamine concentration in proliferating cells is within the range of abundant amino acids like glutamine^2,3^. Polyamine biosynthesis is upregulated in proliferating cells, while its inhibition results in cell cycle arrest^4–6^. In some cases, this arrest resembles terminal differentiation^7–9^. Consistent with proliferating cells requiring high levels of polyamines, ODC1 is a top transcriptional target of the transcription factor MYC^10^, and upregulation of ODC1 is required for the transforming effects of MYC^11,12^. ODC1 is an irreversible enzyme in metazoans, therefore cells cannot normally recover carbon or nitrogen from existing polyamine pools. Thus, proliferating cells invest a significant fraction of critical nutrients (e.g. glutamine and arginine) in the irreversible production of polyamines, yet their biological role is poorly understood.

**Figure 1.**
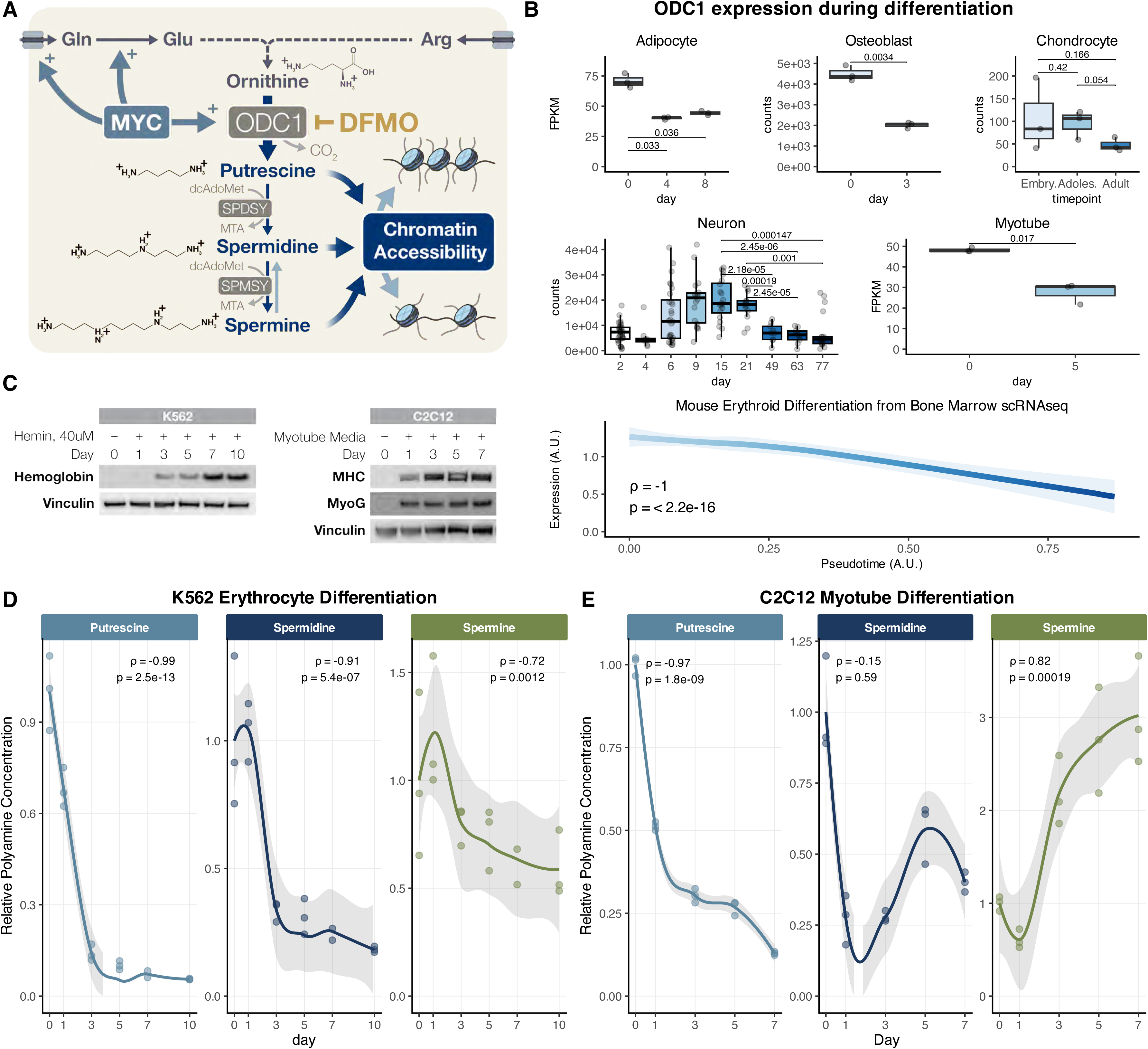
A) Polyamines are synthesized by the irreversible decarboxylation of ornithine by ornithine decarboxylase (ODC1), which is a direct transcriptional target of MYC, and inhibited by the drug difluoromethylornithine (DFMO). B) ODC1 expression from publicly-available RNAseq and scRNAseq (erythroid) data collected over time courses of adipocyte, osteoblast, chondrocyte, neuronal, myotube, and erythroid differentiation. For RNAseq samples, box plots display medians and quartiles of the data. P values correspond to T test with Bonferroni correction. For scRNAseq data, plot shows mean +/- SEM and rho and p value correspond to Spearman test. C) Western blots showing induction of the erythroid marker Hemoglobin in K562 erythroblastic human leukemia cells during ten days of differentiation, and of the myotube markers Myosin Heavy Chain and MyoG in C2C12 cells after seven days of treatment with myotube differentiation media. D) Quantification of relative polyamine levels over a time course of differentiation in K562 cells. Abundance of each polyamine species was normalized to cell number. Spearman Correlation coefficient and P value shown. E) Quantification of relative polyamine levels over a time course of differentiation in C2C12 cells. Abundance of each polyamine species was normalized to cell number. Spearman Correlation coefficient and P value shown.

Mechanistically, polyamines have been shown to perform multiple biological functions, including participating in the post-translational modification of the translation initiation factor eIF5A^13,14^ and regulating the activity of inwardly rectifying potassium channels^15^. Earlier work has also shown that polyamines interact with negatively-charged macromolecules like DNA and RNA. This interaction with DNA can alter chromatin accessibility, nucleosome condensation^16,17^, and increase transcriptional efficiency *in vitro*^16,18–21^, but whether and how this occurs *in vivo* is unknown. Here we show that polyamines are enriched in the nucleus, where their interaction with chromatin enables increased accessibility of the histone tail to chromatin-modifying enzymes. These interactions are critical for the regulation of cell state. By decreasing polyamine levels, differentiating cells limit noisy chromatin dynamics, enabling faithful lineage commitment. Conversely, increasing intracellular polyamine concentrations facilitates lineage plasticity and reprogramming. Thus, polyamines alter the flexibility of chromatin landscapes in an enzyme- and sequence-agnostic manner, facilitating a broad diversity of cell state transitions.

## Results

### Polyamine biosynthesis is decreased during differentiation

Since polyamine content is highest in undifferentiated cells, we wondered if polyamine metabolism is regulated during differentiation. To explore transcriptional regulation of polyamine biosynthesis, we mined published RNAseq datasets in diverse models of differentiation. Across many models (3T3L1 cells during adipocyte differentiation, 10T1/2 cells during osteoblast differentiation, human chondrocyte tissue samples during early development, human embryonic stem cells during neuronal differentiation, C2C12 cells during myotube differentiation, and human hematopoietic cells), we found that ornithine decarboxylase (ODC1) expression decreased as cells underwent fate commitment (Fig. 1B and Fig. S1A-H). Expression of spermidine synthase was also decreased during differentiation, but spermine synthase expression was not clearly correlated with differentiation state in every model (Fig. S1C-H). We next asked if cellular polyamine content similarly decreased as cells differentiated. We used two well established models of differentiation: K562 myelogenous leukemia cells differentiated into erythrocytes, and murine myoblasts (C2C12 cells) differentiated into myotubes. We validated induction of differentiation by western blot for lineage markers (Fig. 1C). To quantify cellular polyamine content, we used an optimized protocol for polyamine extraction and derivatization for gas chromatography mass spectrometry (GC-MS; K562) or LC-MS (C2C12; See Materials and Methods)^22^. In both models, we found that levels of polyamines decreased during differentiation (Fig. 1D and E) and remained low as cells continued differentiating. This was especially the case in K562 cells, where levels of all polyamine species decreased markedly during differentiation (Fig. 1D). For the myoblast to myotube model (C2C12), overall polyamine levels were lower, requiring more sensitive LCMS analysis, and putrescine levels showed a clear sustained decrease during differentiation, though this decrease was not as marked for spermidine and spermine (Fig. 1E). Finally, we mined a published metabolomics data set collected over the first ninety-six hours of zebrafish development and found that putrescine and spermidine levels increased for 48h after fertilization, then decreased as cells began to differentiate (Fig. S1I and Fig. S1J; N.B. spermine abundance was not reported). These data demonstrate that across multiple models and at both the transcriptional and metabolite-levels, polyamine levels decrease during lineage commitment.

### Polyamine depletion induces a differentiated-like cell state

We next asked if polyamine depletion might play a causative role during differentiation. To test this, we depleted cells of polyamines and assessed their differentiation states. We used CRISPR-Cas9 to genetically ablate ODC1 in 10T1/2 cells, a murine mesenchymal progenitor cell line with multipotent potential^23^. We outgrew single cell clones in the presence of exogenous putrescine (Fig. S2A), and validated the effect of ODC1 knockout on polyamine content with GC-MS. The abundance of all three polyamine species was decreased in knockout cells (Fig. 2B, Fig. S2B). We also targeted ODC1 pharmacologically by treating cells with the irreversible ODC1 inhibitor di-fluoro-methyl-ornithine (DFMO), which decreased the abundance of all three polyamine species to a similar extent as ODC1 knockout (Fig. S2C).

**Figure 2.**
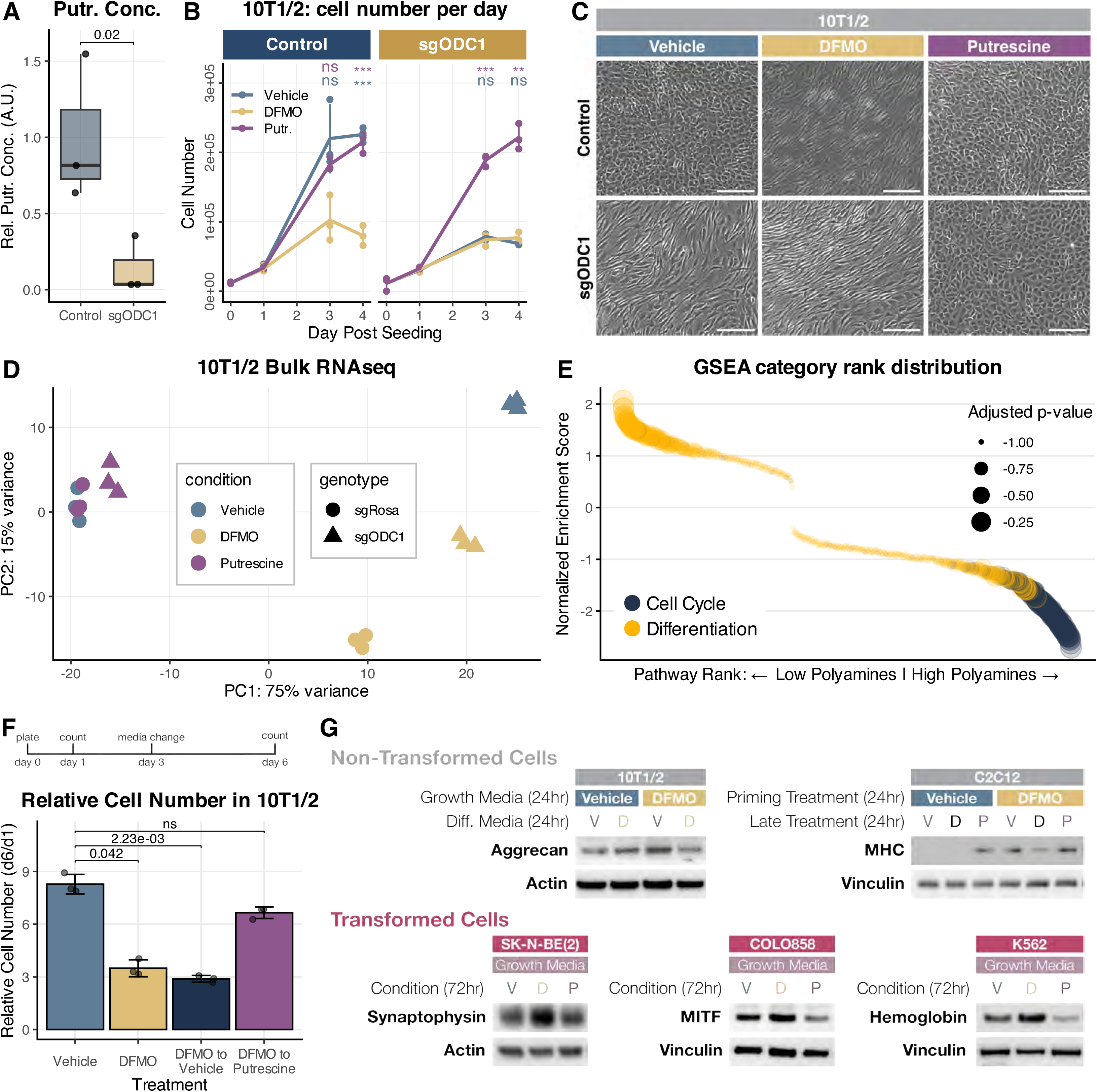
A) Control (sgRosa) and ODC1 knockout (sgODC1) cells were plated in media containing vehicle, 1 mM DFMO, or 1 mM putrescine, and cells were counted one, three, and four days after seeding. Data presented are mean +/- standard error of three biological replicates. P values calculated by T test with Bonferroni correction. B) Relative putrescine concentrations in control (sgRosa) and ODC1 knockout (sgODC1) cells following three days of growth in putrescine-free media. Putrescine abundance was normalized to cell counts. Error bars correspond to standard error. P values calculated by one-tailed t-test. C) Morphology of control (sgRosa) and ODC1 knockout (sgODC1) cells following three days of indicated treatments (1 mM DFMO, 1 mM putrescine). Scale bar is 275 µM. D) Principal component analysis of RNAseq data from control (sgRosa) and ODC1 knockout (sgODC1) cells. RNA was extracted from cells following three days of indicated treatments (1 mM DFMO, 1 mM putrescine). E) Gene set enrichment analysis comparing low-polyamine samples (control cells treated with DFMO, and ODC1 knockout cells treated with vehicle or DFMO) to high-polyamine samples (control cells treated with vehicle or putrescine, and ODC1 knockout cells treated with putrescine). F) 10T1/2 cells were treated with either vehicle or DFMO-containing media and moved into indicated treatments after three days. Cells were counted six days after plating (three days after media change). Data presented are mean +/- standard error of three biological replicates. P values calculated by Dunn’s test. G) Western blots showing markers of differentiation in progenitor and transformed cell lines. 10T1/2 cells were treated with the indicated treatments (1 mM DFMO) in growth media for 24 hours and then moved to differentiation media for 24 hours. C2C12 cells were treated with the indicated treatments (1 mM DFMO, 1 mM putrescine) in differentiation media. All transformed cell lines were treated with 1 mM DFMO in growth media.

Polyamines are known to be required for cell proliferation^4,24^. Consistent with this, DFMO-treated and ODC1 knockout cells failed to proliferate following three days of polyamine depletion (Fig. 2A). We did not observe decreases in cell number or viability (Fig. S2D) which would indicate widespread cell death, suggesting that the effect of polyamine depletion is cytostatic. EdU incorporation and cell cycle analysis by flow cytometry showed that polyamine depleted cells were specifically arrested in G1 (Fig. S2E). We were able to rescue the cytostatic effect of polyamine depletion in ODC1 knockout cells by supplementing media with 0.5 mM putrescine, validating enzymatic loss of function and indicating that knockout cells behaved like putrescine auxotrophs (Fig. 2A).

To our surprise, polyamine depletion resulted in changes in morphology in 10T1/2 cells. While wild-type 10T1/2 cells are normally cuboidal in shape, ODC1 knockout and DFMO-treated cells became elongated and spindly (Fig. 2C). The combination of cell cycle arrest in G1 and changes in morphology suggested that polyamine-depleted cells might be undergoing a differentiation-like change in cell state. To explore whether polyamine depletion was leading to differentiation, we carried out RNAseq on control and ODC1 knockout cells treated with DFMO or exogenous putrescine, grouped the cells according to low or high polyamine content as assessed by GC-MS, and performed differential gene expression analysis (Fig. 2D). Genes upregulated in polyamine-depleted samples were involved in the regulation of diverse cell differentiation outcomes, multicellular development, and intracellular signal transduction (Fig. 2E, top terms shown in Fig. S2F). Conversely, gene sets down-regulated in polyamine-depleted cells were involved in cell cycle regulation (Fig. 2E, top terms shown in Fig. S2G), agreeing with the cytostatic effect of polyamine depletion. Critically, polyamine depletion did not lead to upregulation of genes associated with any one lineage, suggesting that polyamine depletion is not inducing differentiation on its own, but rather promoting a “differentiation-like” or “differentiation-primed” cell state.

A common feature of terminal differentiation is irreversible exit from the cell cycle. To explore whether the arrest in G1 observed in polyamine-depleted cells was reversible, we treated cells with DFMO for three days, then allowed them to recover for three days in media without drug. Polyamine-depleted cells did not resume proliferation after drug washout (Fig. 2F). Thus, DFMO treatment induces a stable cell state change which is not reversed when ODC1 function is restored. Taken together, these data suggest that polyamine depletion pushes cells towards a more differentiated phenotype.

To test whether polyamine depletion induces differentiation in other contexts, we treated a panel of cell lines with DFMO and assessed expression of lineage commitment markers by western blot. In 10T1/2 cells cultured in chondrocyte differentiation media, DFMO treatment either 24 hours before or at the onset of differentiation increased differentiation marker expression (Fig. 2G). However, DFMO had a negative effect on differentiation if present once the differentiation process was underway. The multipotent nature of c10T1/2 cells enabled us to probe whether differentiation into other lineages was similarly enhanced by polyamine depletion. As opposed to enhancing chondrocyte differentiation, polyamine depletion with DFMO impaired adipocyte differentiation (Fig. S2H). Polyamine depletion was also able to enhance differentiation of C2C12 myoblasts into myotubes; DFMO treatment during the myoblast stage was able to prime differentiation, but impaired differentiation if added during the final stages (Fig. 2G). As opposed to these non-transformed models of differentiation, in three cancer cell lines tested (SK-N-BE(2) neuroblastoma, COLO858 melanoma, and K562 acute myeloid leukemia) polyamine depletion resulted in decreased proliferation (Fig. S3A) and up-regulation of markers of differentiation (Fig. 2G). Thus, when polyamines are depleted, non-transformed progenitors are primed for increased differentiation towards a defined lineage and, in the case of a multipotent progenitor, away from an alternative lineage as seen for adipocytes. Conversely, upon polyamine depletion, transformed cells are pushed towards a differentiation-like state defined by their cell of origin. These data suggest that polyamine depletion is not merely a consequence of lineage commitment, as experimental depletion is sufficient to drive cells towards a more differentiated phenotype.

Polyamines are known to contribute to translational homeostasis through spermidine’s role as a substrate for eIF5a hypusination^13,14^. Hypusinated eIF5a facilitates the translation elongation and termination of polyproline-rich and tripeptide repeat-containing proteins^25–27^, and it has been suggested that polyamines impact growth and lineage fidelity through their regulation of hypusination^28^. To investigate whether polyamine-depleted 10T1/2 cells were undergoing proteotoxic stress as a result of impaired translation, we pulsed cells with the puromycin analogue O-propargyl-puromycin (OPP) and quantified global translation by flow cytometry. We did not observe changes in bulk translation following depletion or addition of polyamines (Fig. S3B-C), nor did we observe induction of phosphorylated eIF2a, a readout of the integrated stress response (Fig. S3D)^29^. To explore the impact of hypusination on differentiation, we pharmacologically inhibited deoxyhypusine synthase (DHPS), the enzyme responsible for forming the first intermediate deoxyhypusine residue on eIF5a. We validated decreased hypusine levels following DHPS inhibition by western blot. Markers of differentiation either decreased or did not change (Fig. S3E) after DHPS inhibition. These data suggest that the differentiation phenotypes observed after polyamine depletion are not due to impaired hypusination.

### Polyamines are enriched in the nucleus

Subcellular compartmentalization of metabolites can restrict the effects of small molecules to specific parts of a cell, even in the absence of membrane-bound organelles^30,31^. We reasoned that identifying where polyamines were concentrated in cells might help illuminate their mechanism of action. Resolving subcellular localization of metabolites by mass spectrometry is limited by the spatial resolution of MALDI (Matrix-Assisted Laser Desorption/Ionization)^32^, so we set out to identify an alternative imaging-based approach. We synthesized a clickable putrescine probe consisting of putrescine with an azide group that can be covalently bound to a fluorophore-alkyne label *in vitro* or *in vivo* (Fig. 3A). We treated live cells with the un-clicked probe, fixed, permeabilized and performed click-chemistry to a fluorophore alkyne, and then imaged cells using confocal microscopy. We did not observe any fluorescence signal when cells were exposed to the probe without an alkyne fluorophore (Fig. S4A). Imaging after clicking the probe to a fluorophore revealed stark nuclear localization in 10T1/2 cells (Fig. 3B). DFMO treatment of cells prior to incubation in polyamine probe increased probe signal, presumably due to upregulation of polyamine transporters^33–35^. Similar effects were seen in ODC1 knockout 10T1/2 cells when grown in the absence of exogenous putrescine (Fig. S4B). Nuclear probe localization was also observed with spermidine and spermine probes (Fig. 3C and Fig. 3D), and in RPE-1 and U-2 OS cells (Fig. S4D and Fig. S4E). Critically, incubation in D-alanine and L-ornithine azide probes did not reveal nuclear localization, confirming that nuclear localization is a property unique to polyamines (Fig. S4A). To determine if polyamines are primarily associated with chromatin (DNA and protein), or RNA species, we incubated cells in the putrescine probe, and then treated cells with RNAse. Following RNAse treatment, cells were washed thoroughly and imaged. Nuclear probe localization was observed regardless of RNAse treatment (Fig. S4F, suggesting that polyamines are primarily associated with chromatin in the nucleus. To validate our localization data *in vivo*, we manually dechorionated zebrafish embryos at the four-cell stage and exposed them to the putrescine probe for two hours, then fixed, permeabilized, and performed click chemistry. The putrescine probe again concentrated in nuclei, especially at the nuclear periphery (Fig. S4G). Taken together, these data suggest that all three polyamine species are enriched in the nucleus, where they primarily associate with chromatin.

**Figure 3.**
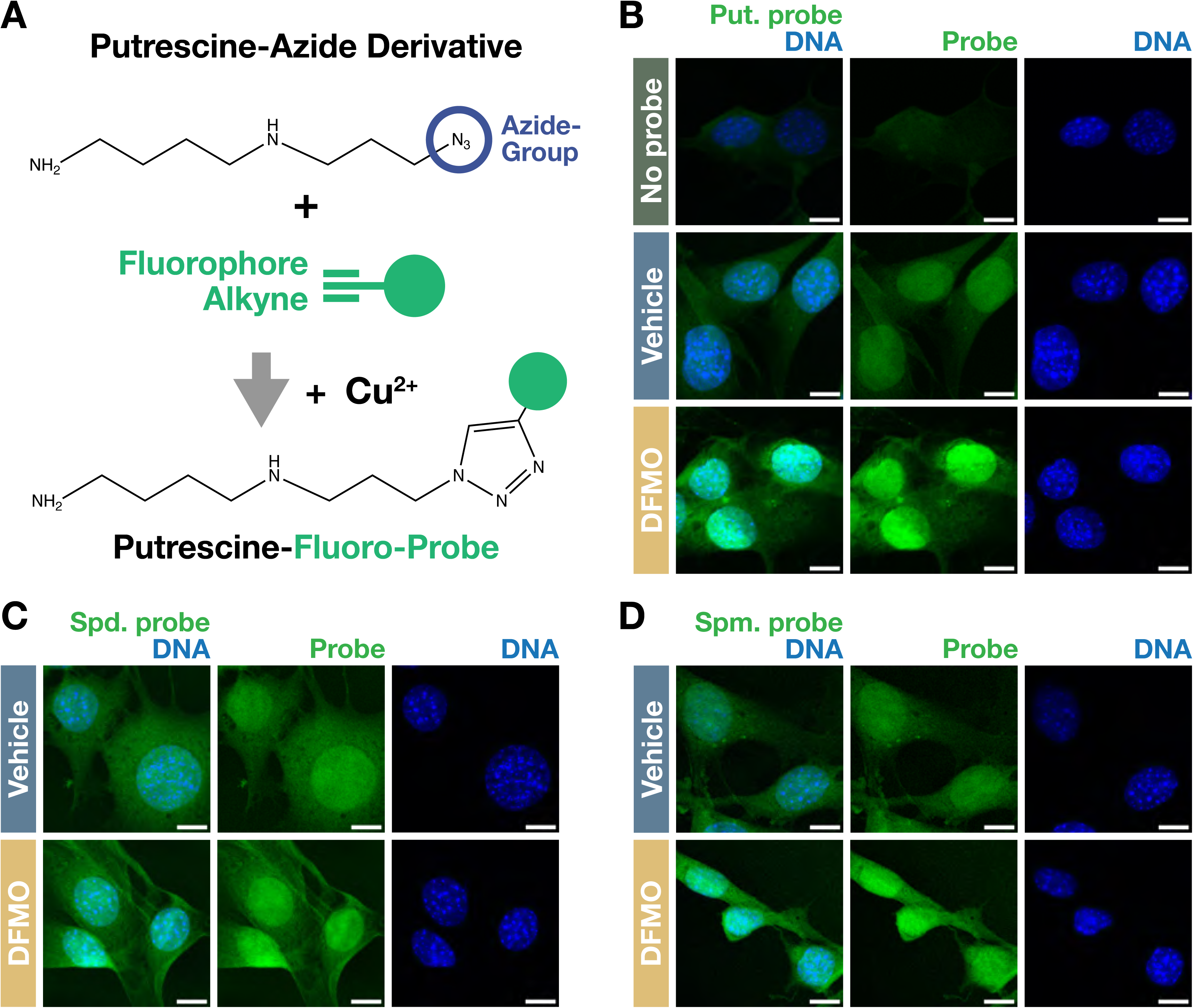
A) Structure of the putrescine-azide probe. B) Intracellular localization of the putrescine probe in 10T1/2 cells visualized by confocal microscopy following three days of the indicated treatments (1 mM DFMO, 1 mM putrescine). Scale bars are 5 µM. C) Intracellular localization of the spermidine probe in 10T1/2 cells visualized by confocal microscopy following treatments as per B). Scale bars are 5 µM. D) Intracellular localization of the spermine probe in 10T1/2 cells visualized by confocal microscopy following treatments as per B). Scale bars are 5 µM.

### Polyamines increase the activity of diverse histone-modifying enzymes

Polyamines have been reported to alter DNA/DNA and DNA/protein interactions^36,37^. Given their nuclear localization and the effects of polyamine depletion on differentiation, we set out to explore the impact of polyamine depletion on chromatin. We first profiled chromatin accessibility broadly by performing micrococcal nuclease (MNase) digestions on wildtype and ODC1 knockout 10T1/2 cells and quantifying the abundance of mononucleosome fragments. Mononucleosome abundance was increased in ODC1 knockout cells (Fig. S5A), suggesting that global chromatin accessibility increases upon polyamine depletion. To determine the specificity of this change in chromatin accessibility, we performed ATACseq (Assay for Transposase-Accessible Chromatin using sequencing) on 10T1/2 cells treated with DFMO or putrescine (Fig. 4A and Fig. S5B). Addition of putrescine had minimal effects on chromatin accessibility, but polyamine depletion resulted in thousands of regions of chromatin with either increased or decreased accessibility (Fig. 4A, Fig. S5C, samples normalized to E. coli spike-in DNA). As we had seen with MNAse treatment, and consistent with earlier reports^21^, we noticed an overall increase in chromatin accessibility in DFMO-treated samples. At a more granular level, genes with increased accessibility in transcriptional start sites following polyamine depletion were associated with the regulation of anatomical structure size, morphogenesis, and cell adhesion (Fig. S5D), and de novo motif enrichment analysis revealed binding sites for transcription factors known to coordinate lineage and responses to diverse stressors (Fig. S5E). Together with our transcriptomic data, these results suggest that polyamine depletion initiates a concerted rewiring of epigenetic and transcriptional landscapes, pushing cells towards a more differentiated state.

**Figure 4.**
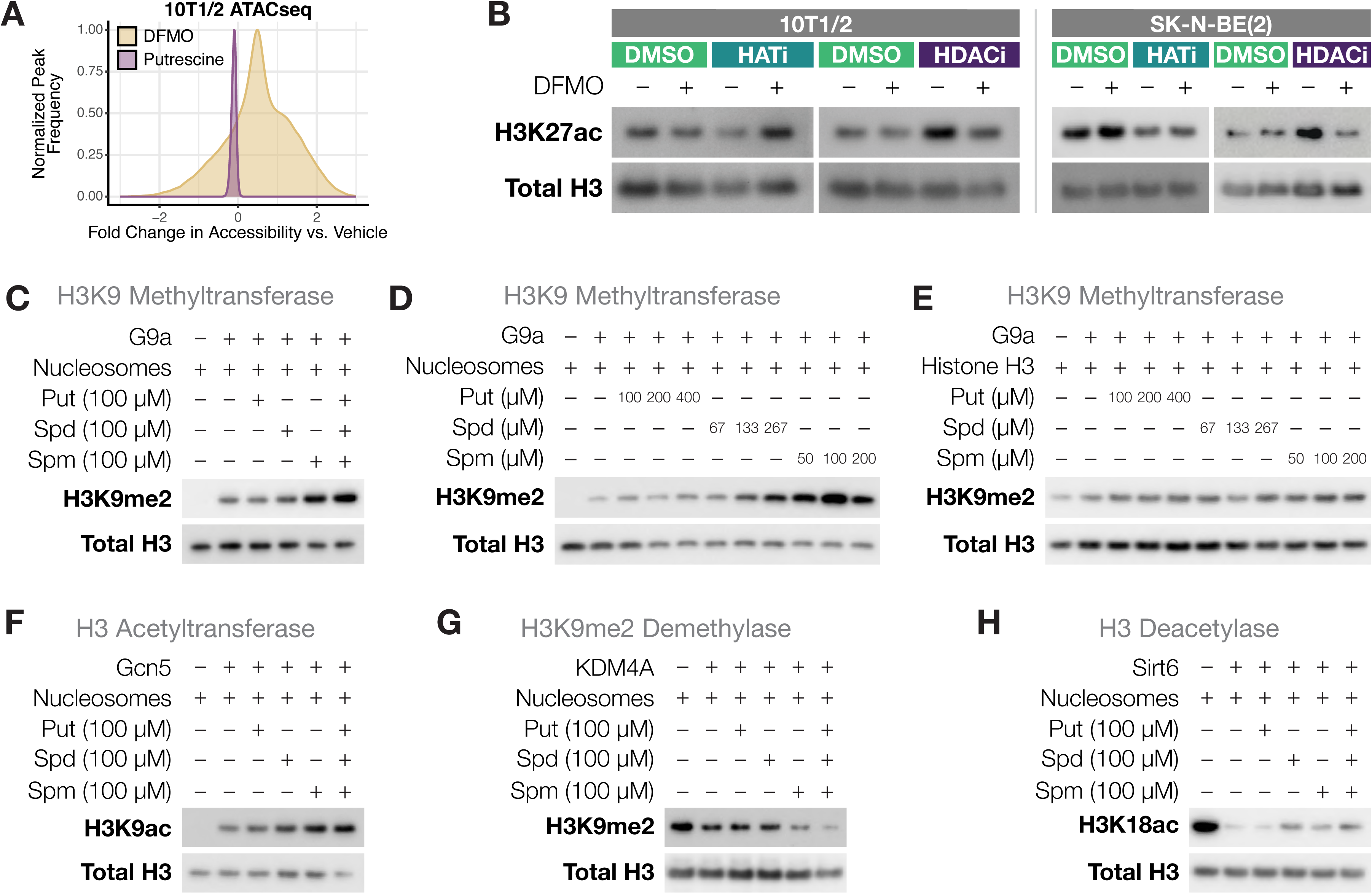
A) Distribution of the fold change in peak accessibility in 10T1/2 cells treated with 1 mM DFMO or 1 mM putrescine versus vehicle for three days as determined by ATACseq. B) Western blots showing abundance of H3K27ac following treatment with a histone acetyltransferase inhibitor (25 µM JG2016) or histone deacetylase inhibitor (0.5 µM Vorinostat) in the presence or absence of 1 mM DFMO. Cells (10T1/2 or SK-N-BE(2)) were pre-treated with DFMO for 24 hours and then transferred into media with HAT and HDAC inhibitors with or without DFMO for 24 hours. C) Western blot showing H3K9me2 abundance following *in vitro* methyl-transferase reaction by G9a. Recombinant G9a was incubated in the presence of recombinant K9me2-modified mononucleosomes and 100 µM putrescine, spermidine, or spermine for one hour at 37°C. Reaction was quenched with SDS, boiled, and assessed by western blot. D) *In vitro* methyl-transferase reactions with G9a and nucleosome as per C) incubated in charge-matched concentrations of putrescine, spermidine and spermine. E) *In vitro* methyl-transferase reactions with G9a and histone H3 incubated in charge-matched concentrations of putrescine, spermidine and spermine. F) Western blot showing H3K9ac abundance following *in vitro* acetyl-transferase reaction with recombinant Gcn5, mononucleosomes and indicated polyamine species. Reactions were incubated for one hour at 37°C, and quenched with SDS. G) Western blot showing H3K9me2 abundance following *in vitro* demethylation reaction. Recombinant Kdm4a was incubated in the presence of recombinant H3K9me2 mononucleosomes and 100 µM putrescine, spermidine, or spermine for two hours at 37°C. H) Western blot showing H318ac abundance following *in vitro* deacetylation reactions. Recombinant Sirt6 was incubated with recombinant acetylated nucleosomes and 100 µM putrescine, spermidine, or spermine for one hour at 37°C.

Changes in chromatin accessibility are coordinated in part by enzymes that deposit modifications on histone tails. To characterize histone tail modifications following polyamine depletion, we assessed the levels of a panel of histone modifications by western blot, finding that some modifications increased, some decreased, and some were unaltered (Fig. S5F). These changes were not due to the cytostatic effects of polyamine depletion, as arresting cells in G1 with palbociclib did not induce changes in H3K18ac or H3K9me2 (Fig. S5G). Additionally, we did not observe changes in global DNA 5mC content (Fig. S5H). We next wanted to understand how polyamine depletion might alter histone tail modifications. A challenge of interpreting changes to bulk histone modifications is that they are the sum of the activities of both writer and eraser enzymes. Therefore, we set out tease apart the effect of polyamine depletion on different classes of histone-modifying enzymes. We exposed 10T1/2 and SK-N-BE(2) cells to inhibitors of histone acetyltransferases (JG2016, HATi) or histone deacetylases (Vorinostat, HDACi) and depleted polyamines with DFMO. We reasoned that DFMO treatment in the context of histone writer inhibition (HATi) would allow us to assess the impact of polyamine depletion on histone deacetylase activity. HATi treatment decreased histone acetylation, reflecting the accumulated activity of deacetylases (Fig. 4B). Polyamine depletion in the presence of histone acetyltransferase inhibition attenuated this decrease, suggesting that polyamine depletion decreases the activity of histone deacetylase enzymes. The same effect was seen after inhibition of deacetylases – increased acetylation following HDACi treatment was decreased by concomitant polyamine depletion – as well as methyltransferase and demethylase inhibition, to varying extents in the two cell lines tested (non-transformed 10T1/2 cells and transformed SK-NE-B(2) cells) (Fig. S5I). This data suggests that polyamine depletion decreases the activity of diverse classes of histone-modifying enzymes.

To directly query the impact of polyamines on histone-modifying enzymes, we performed *in vitro* histone modification reactions with purified histone-modifying enzymes, cognate substrates, recombinant nucleosomes, and polyamines. These assays have successfully demonstrated the role of metabolites in histone-modifying enzyme activity^38^. As we observed decreased activity of histone methyltransferases, acetyltransferases, demethylases and deacetylases in polyamine depleted cells, we performed *in vitro* reactions with members of each of these four distinct enzyme classes in the presence of increasing polyamines, using the histone modifications detected by western blotting as a read-out of catalytic activity. Spermine and, to a lesser extent, spermidine increased the catalytic activity of the H3K9 histone-methyltransferase (HMT) G9a on nucleosome substrates (Fig. 4C) while putrescine had little effect. Given the electrostatic charge relationship of the different species (putrescine, 2+; spermidine, 3+; spermine, 4+), we wondered whether the increased efficiency of spermine was related to its higher charge content. However, even with concentrations of putrescine and spermidine that matched the charge concentration of spermine, spermine had the largest effect on increasing G9a activity (Fig. 4D). To determine whether the effect of spermidine and spermine on G9a methyl-transferase activity was dependent on an interaction of the polyamine with the nucleosome substrate or the enzyme itself, we repeated the methyl-transferase reaction with purified histone H3. In this case, none of the polyamines had any effect on catalytic activity (Fig. 4E). These results show that spermine, and to a lesser extent, spermidine, can increase the ability of G9a to methylate histone H3 in the context of the nucleosome.

We next turned to evaluating the effects of polyamines on other histone-modifying enzymes. Spermine and spermidine were also able to increase the efficiency of the histone acetyl-transferase (HAT) GCN5 on nucleosome substrates (Fig. 4F). This effect was independent of charge (Fig. S5J) and was not seen when histone H3, as opposed to intact nucleosomes, was provided as a substrate (Fig. S5K). To evaluate the effect of polyamines on histone demethylation (KDM) activity, we incubated recombinant KDM4A, a Histone H3 K9me2/3 demethylase, with recombinant H3K9me3-modified nucleosomes. Again, spermine increased the efficiency of H3K9me2/3 demethylation by KDM4A (Fig. 4G), in agreement with our *in vivo* results. Neither putrescine nor spermidine increased the efficiency of this reaction. As shown for the histone writers G9a and GCN5, the polyamine-specific increase in catalytic activity required intact nucleosomes, as they were not seen with purified histone H3 (Fig. S5L). Finally, we evaluated the effect of polyamines on the catalytic activity of the histone deacetylase SIRT6. In this case, neither individual polyamines nor a pool of polyamines was able to increase the efficiency of acetylated H3 deacetylation in nucleosomes (Fig. 4H) or free histone H3 (Fig. S5M). These data suggest that spermine, to a lesser extent spermidine, and a pool of polyamines, can increase the catalytic activity of multiple subclasses of histone-modifying enzymes on nucleosome substrates but not free histone protein.

### Polyamines increase the accessibility of the histone tail in the context of the nucleosome

Deuterium exchange mass spectrometry has shown that histone tails are solvent protected, likely due to their electrostatic interactions with the DNA backbone^39^. This interaction has also been observed by NMR analysis^40–42^. To serve as substrates of enzymatic activity, polycationic histone tails must dissociate from negatively charged nucleosomal gyre DNA. We hypothesized that this dissociation was favored by polyamines, which would bind electrostatically to the DNA backbone and allow the histone tail to become freely available to diverse classes of histone-modifying enzymes. This model would explain why in our *in vitro* catalytic assays, polyamines only improved enzymatic activity when intact nucleosomes, rather than free histone H3, were the substrate. Importantly, we did not find evidence that polyamines at the concentrations used in our *in vitro* assays could denature DNA from the nucleosome core (Fig. S5N)^43^. Another prediction of this model is that as opposed to modifications of the relatively unstructured histone tail, modifications to the core histone particle, like histone H3K79 di-methylation catalyzed by the enzyme DOT1L, would not be impacted by polyamines. To test this, we treated 10T1/2 cells with DFMO alone or in combination with EPZ5676, an inhibitor of DOT1L^44^, and found that H3K79me2 levels were unchanged (Fig. S5O). We also performed *in vitro* histone methylation reactions with recombinant DOT1L protein and found that enzymatic activity was unaffected by addition of polyamines (Fig. S5P).

To further characterize how polyamines affect tail accessibility in nucleosomes, we performed *in vitro* trypsin digestion assays^45^. In a time course of nucleosome digestion in vehicle- and spermidine-incubated reactions, spermidine increased the ability of trypsin to degrade the largest molecular weight band corresponding to intact H3 histone (Fig. 5A and Fig. S6A). Spermine was also able to increase the ability of trypsin to digest nucleosomal substrates, though in this case efficiency was more clearly demonstrated in the disappearance of intermediate digestion products (Fig. 5B and Fig. S6A). A combination of all three natural polyamines was similarly able to increase the efficiency of trypsin digestion of nucleosomal histones (Fig. S6B). As we had seen for the catalytic assays with chromatin-modifying enzymes, none of the polyamines had any effect on the ability of trypsin to digest recombinant free histone H3 (Fig. S6C). These results support a model whereby polyamines increase the accessibility of the histone tail in the context of an intact nucleosome.

**Figure 5.**
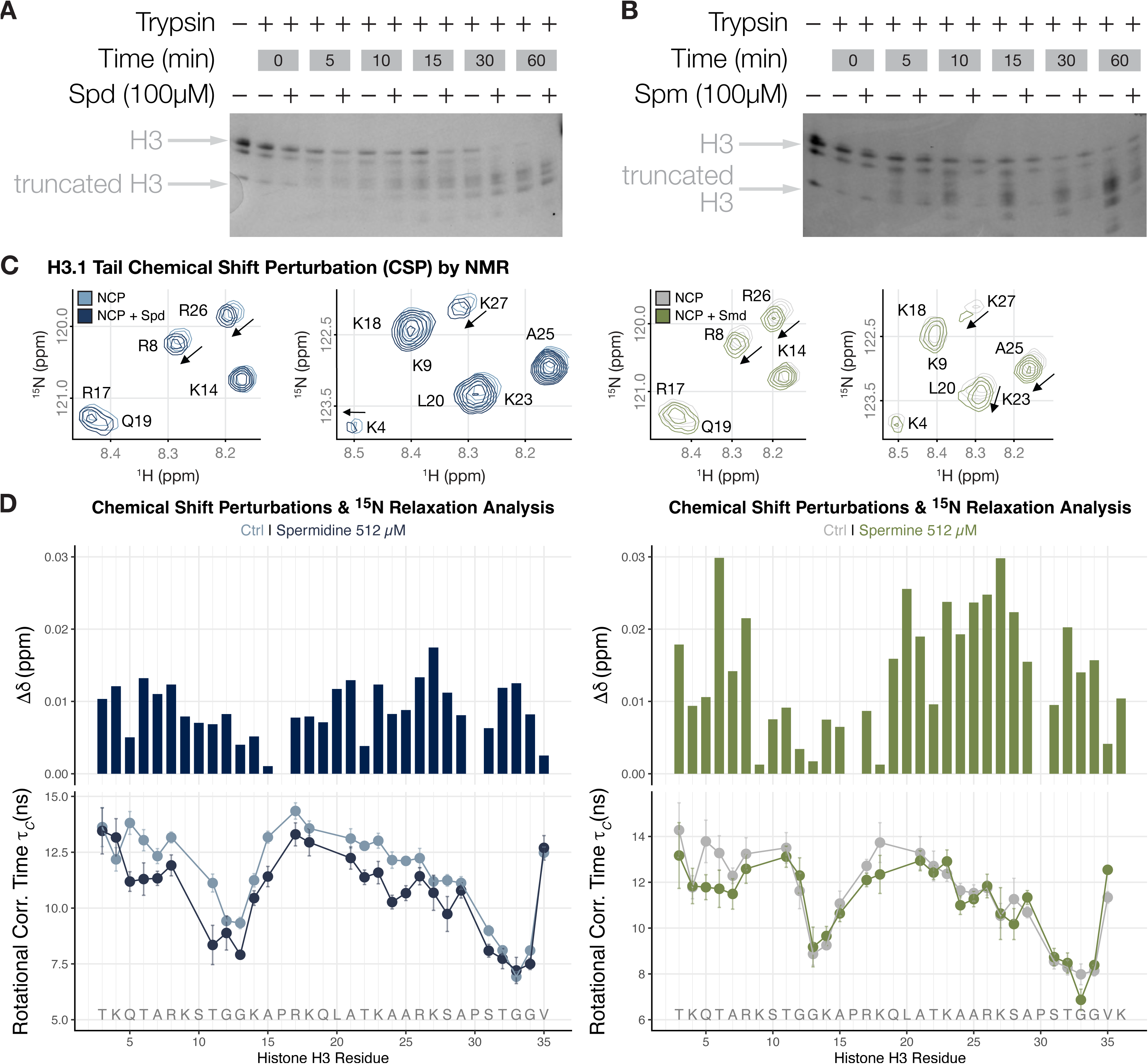
A) Time-course of trypsin-digestion of nucleosome substrate in the presence or absence of 100 µM spermidine visualized by Imperial stain of SDS-page-separated samples. B) Time-course of trypsin-digestion of nucleosome substrate in the presence or absence of 100 µM spermine visualized by Imperial stain of SDS-page-separated samples. C) Overlay of the selected regions in the ^1^H,^15^N-HSQC spectra of ^15^N-H3-NCP (128 µM) with (dark blue) and without (light blue) 512 µM spermidine trichloride (left) and with (green) and without (grey) 512 µM spermine trichloride. D) Top) Normalized chemical shift difference as a function of H3 tail residues in ^15^N-H3-NCP at 512 µM spermidine (left) or spermine (right) trichloride concentration. The dotted line represents the average shift. Bottom) Rotational correlation time (*τ*_C_) of H3 tail residues in ^15^N-H3-NCP with (dark blue/green) and without (dark blue/green) spermidine/spermine.

We next wanted to probe the structural changes that polyamines have on histone tail conformational dynamics, as suggested by the *in vitro* enzymatic and trypsin tail digestion assays. Previous NMR studies have shown that H3 tails are associated with DNA in the context of nucleosomes, forming what is termed a fuzzy complex that is inhibitory to binding and can be regulated by histone post-translational modifications and nucleosome composition^41,42,45–47^. To test if spermidine can alter the conformational ensemble of the H3 tail, we performed chemical shift perturbation (CSP) experiments on a nucleosome core particle (NCP) in which H3 was ^15^N-labelled (^15^N-H3-NCP) (Fig. S6D-E). With this labeling scheme, H3 tails are observable within the context of the nucleosome. We collected ^1^H,^15^N heteronuclear single quantum coherence (^1^H,^15^N-HSQC) spectra on the ^15^N-H3-NCP in the absence and presence of spermidine (Fig. 5C-D and Fig. S6F). In the presence of spermidine we observed CSPs consistent with an altered H3 tail conformational ensemble. These CSPs were observable across the entire H3 tail, indicating that all residues are affected by spermidine. Notably, the direction of the CSPs is consistent with previously observed CSPs seen upon addition of increasing monovalent or divalent salt concentrations^42^ which are known to enhance the accessibility of the H3 tail. We also carried out chemical shift perturbations on a nucleosome core particle in the absence and presence of spermine (Fig. 5C-D and Fig. S6F). In this case, we observed greater CSPs overall as compared to spermidine, but also saw these concentrated in two regions: a smaller region closer to the N-terminus comprising H3 residues 3-8 and a larger region encompassing H3 residues 19-29. To assess how spermine and spermidine alter the conformational dynamics of the H3 tails we carried out ^15^N-relaxation experiments and calculated the amino acid specific correlation times (tC), which report on the overall mobility of each amino acid. Notably, spermidine led to a consistent decrease in tC for residues 5-29, indicating an increase in the conformational dynamics of the majority of the H3 tail. While spermine also led to an increase in conformational dynamics, the effects were more localized, affecting residues 3-11 and 15-21. Together, these data reveal that spermine and spermidine alter both the conformational ensemble and dynamics of the H3 tails but in a manner specific to the polyamine. These *in vitro* experiments provide orthogonal support for a model where polyamines change the conformational ensemble of the H3 tail, likely by directly associating with the nucleosomal DNA to free and increase tail accessibility.

### Exogenous polyamines facilitate reprogramming

We have so far shown that polyamine depletion induces a lineage primed cell state by altering the ability of histone-modifying enzymes to access histone tails. We wondered if exogenous polyamines might increase chromatin dynamics and facilitate reprogramming of terminally differentiated cells. Supporting this, though cells pre-treated with DFMO for three days did not re-initiate proliferation after removal of drug, they did proliferate if media was supplemented with excess putrescine (Fig. 2F). ODC1 is a direct transcriptional target of c-Myc, one of the four Yamanaka reprogramming factors^48^. We hypothesized that the role of c-Myc during reprogramming might be in part to increase polyamine biosynthesis, thereby facilitating the chromatin remodeling necessary for de-differentiation. To test this, we reprogrammed mouse embryonic fibroblasts expressing an Oct4-GFP reporter by doxycycline-inducible expression of either three (Oct4, Sox2, and Klf4; OSK) or four (Oct4, Sox2, Klf4, and Myc; OSKM) Yamanaka reprogramming factors for 12 days in the presence of DFMO or putrescine. We assayed stable reprogramming efficiency by withdrawing cells from doxycycline for four days, then quantifying Oct4-GFP expression and colony number (Fig. 5A-B). Reprogramming with OSK resulted in low reprogramming efficiency with a small population of Oct4-expressing cells, while OSKM expression led to reprogramming efficiencies comparable to previous reports^49,50^. Treatment of MYC-less OSK-expressing cells with putrescine increased Oct4-GFP reporter expression and reprogramming efficiency nearly to the levels of OSKM-reprogrammed fibroblasts (Fig. 5B-D), while DFMO treatment ablated reprogramming capacity in OSK- or OSKM-expressing cells. These data show that a primary role of c-Myc during reprogramming is its ability to increase polyamine biosynthesis.

### Exogenous polyamines can rescue the effects of losing histone tail acetylation

To determine whether the phenotypic consequences of increasing or decreasing polyamines were mediated by their effects on histone tail accessibility, we focused on histone acetylation. Histone tail acetylation has been thought to mediate effects on chromatin function and gene expression by increasing tail accessibility^41,51–53^. Consistent with this, reprogramming from pluripotency is enhanced by histone deacetylase inhibitors^54,55^ and inhibition of histone acetylation inhibits reprogramming^56^. Given that inhibiting HATs reduces histone tail accessibility, we reasoned that if exogenous polyamines could rescue phenotypes that result from acetyl-transferase inhibition, it would suggest that polyamines exert their effects primarily through enhancing histone tail accessibility, rather than through alternative mechanisms (e.g. protein translation or chromatin condensation). We repeated our reprogramming assays with inducible four-factor OKSM reprogrammable mouse embryonic fibroblasts and treated cells with a histone acetyl-transferase inhibitor during the 12 day reprogramming period. Inhibition of HAT activity led to almost complete inhibition of Oct4-GFP positive colonies and addition of exogenous putrescine rescued the inhibitory effects of HAT inhibition to levels equivalent to the vehicle-treated condition (Fig. 6E and sFig. 7A). Exogenous putrescine was also able to rescue the inhibitory effect of HAT inhibition on differentiation of 10T1/2 cells to adipocytes, which unlike chondrocyte differentiation, requires polyamines (sFig. 7B). These experiments support a model in which polyamine perturbations drive changes in cell fate as a result of their ability to increase histone tail dynamics (Fig. 6F).

**Figure 6.**
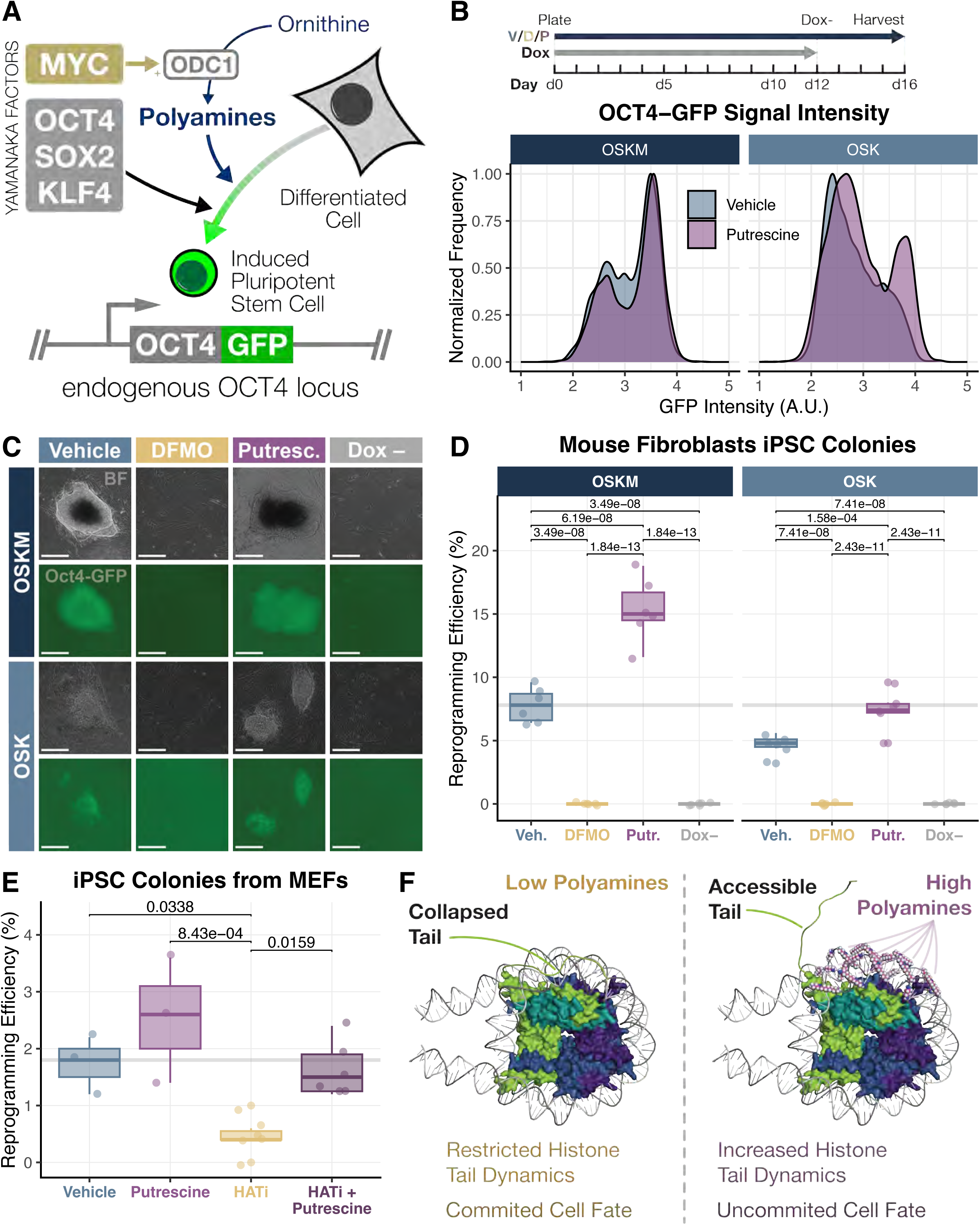
A) Mouse embryonic fibroblasts expressing GFP under the control of the endogenous Oct4 locus were reprogrammed with either three (Oct4, Sox2, Klf4; OSK), or four (Oct4, Sox2, Klf4, Myc; OSKM) Yamanaka factors. B) Top: timeline of reprogramming experiments: MEFs were plated on a feeder layer and treated with doxycycline to induce OSK and OSKM expression. Doxycycline was removed at day 12 and doxycycline-independent colonies assessed after four days. Bottom: Oct4-GFP expression quantified by flow cytometry of indicated cells. C) Morphology and Oct4-GFP expression of reprogrammed colonies following the indicated treatments. Scale bars are 275 µM. D) Reprogramming efficiency quantified by calculating the number of Oct4-GFP-positive colonies following indicated treatments as a percentage of initial cells plated. Each data point represents one biological replicate. P values correspond to Tukey’s test. E) Reprogramming efficiency of OSKM cells treated with indicated compounds and quantified as per D). P values correspond to Tukey’s test. HATi: histone acetyl-transferase inhibitor A-485 0.5 µM. F) Model of effects of polyamines on accessibility of histone tail (H3 tail highlighted).

## Discussion

Here we propose that cells modulate their ability to rearrange chromatin structure by changing polyamine levels. Mechanistically, our data are consistent with a model whereby interactions between histone tails and nucleosomal DNA limit the activity of chromatin-modifying enzymes. Polyamines, by neutralizing negative charges on the DNA backbone, weaken this interaction, allowing increased tail accessibility that facilitates chromatin-modifying enzyme activity (Fig. 6F). This model is supported *in vivo* by the enrichment of polyamines in the nucleus and the fact that DFMO impairs the activity of both chromatin writers and erasers (Fig. 4B and sFig. 5I). Moreover, histone tail dynamics are enhanced by polyamines *in vitro* in 3 orthogonal assays: chromatin-modifying assays, trypsin digestion of histone tails and NMR CSP and relaxation experiments. The model is additionally supported by a) the lack of effects of polyamines when chromatin-modifying enzymes act on histone protein alone (Fig. 4E and sFig. 5K,M); b) that a histone modification that is not on the tail, but on the nucleosome core (H3K79 di-methylation), is unaffected by polyamine levels both *in vivo* and *in vitro* (Fig. S5O-P); and c) that incubation of nucleosomes with polyamines does not denature the nucleosome (Fig. S5N). This histone tail dynamics model provides an explanation for the longstanding observation that spermidine and spermine enhance the efficacy of chromatin-based assays^57^. Our work also underscores the importance of considering these abundant charged metabolites when modeling the structures of protein-protein and protein-nucleic acid interactions. Interestingly, the one chromatin-modifying enzyme reaction that was unaffected by polyamines was histone deacetylation (Fig. 4H). Since acetylated histone tails already show increased dynamics^41,51–53^, it is not surprising that deacetylation efficiency would not be further improved by polyamines.

A question raised by our *in vitro* work is why different polyamine species have different effects on chromatin-modifying enzyme activity and histone tail accessibility. Spermine showed the largest ability to increase tail accessibility while spermidine had slightly less of an effect and putrescine showed no discernible alteration compared to vehicle. However, the effects of spermine on chromatin-modifying enzyme activity were not matched by concentrations of putrescine or spermidine that would provide equivalent charge concentrations. We did find that spermidine readily improved trypsin tail digestion of the H3 N-terminus, while spermine increased digestion of intermediate H3 fragments. This may be explained by the NMR relaxation analysis which suggests that, while spermidine has a broader effect throughout the length of the tail, spermine is more specific, effectively increasing dynamics in two areas centered around the most N-terminal residues and residues 19 to 29 (Fig. 6). More in-depth studies and computational modeling will be required to determine how these different species interact with nucleosomal DNA and differentially change the accessibility of histone tails. However, we know of no situation *in vivo* where polyamine species are present in isolation. Since a mix of polyamines leads to maximal effects on histone tail dynamics, we interpret that these changes also correspond to the changes occurring in nuclear chromatin *in vivo*.

We show that putrescine can substitute for MYC in the setting of reprogramming to pluripotency. Our model suggests that polyamines increase the efficiency of the chromatin remodeling reactions necessary for de-differentiation. Whether this increased efficiency is genome-wide, mediated through transcription factor-dependent recruitment of chromatin-modifying enzymes or through sub-nuclear compartment-driven localization of these metabolites is the focus of ongoing work. Transformed cancer cells also take on a less differentiated state and, in many cases, upregulate polyamine biosynthesis. Accordingly, we found that depletion of polyamines led to decreased proliferation and upregulation of differentiation markers across a panel of cancer cell lines. One intriguing hypothesis is that by upregulating polyamine biosynthesis, transformed cells might acquire the capacity to alter chromatin structure, increasing lineage plasticity and allowing them to better adapt to diverse stressors and environments.

Ascribing the cell fate phenotypes seen when polyamine levels are perturbed *in vivo* to their effects on chromatin structure is made difficult by their abundance and diverse functions. Two experiments support the idea that the cell fate changes seen by perturbing polyamines result from their effects on chromatin. Firstly, one role of polyamines is that spermidine acts as a substrate for the post-translational modification of eIF5A on lysine 50^14,58^. This modification, hypusination, enables translation elongation and termination of mRNAs encoding poly-proline tracts and other tri-peptide motifs^26,27,59^. Previous studies have suggested that polyamine levels impact differentiation through their regulation of hypusination^60^. However, that inhibition of hypusination leads to impaired differentiation of the lines studied in this work, while DFMO and ODC1 loss promote a differentiation-like phenotype, argue that inhibition of hypusination is not mediating the cell state changes seen when polyamines are depleted. Second, we show that exogenous putrescine can rescue the loss of reprogramming or adipocyte differentiation caused by histone-acetyl-transferase inhibition. While it is formally possible that this rescue is mediated by hypusination or other functions of polyamines, histone acetylation increases tail accessibility and thus we interpret that exogenous putrescine is reinstating this effect when acetylation is absent. Thus, we conclude that fluctuations in polyamine levels enable cells to restrict or alter cell states by regulating the accessibility of histone tails.

## Methods and Materials

### Cell culture and drug treatments

10T1/2 and C2C12 cells were cultured in DMEM with 10% FBS and 1% pen-strep. CRISPR knockout lines were generated by cloning pLentiCRISPRv2 (Addgene 98290) plasmids with sgRNA sequences targeting ODC1 or the Rosa safe harbor locus. 293T cells were transfected with PEI with the targeting plasmid and pMD2 (Addgene 12259) and psPAX2 (Addgene 12260) vectors to generate viral supernatants, which was harvested 24-48 hours after transfection and applied to target cells derived from single cell clones in the presence of 8 µg/mL polybrene. Knockout lines were generated from single cells clones after puromycin selection and passaged in media supplemented with 0.5 mM putrescine to rescue growth, then moved to media without putrescine for the duration of experiments. SK-N-BE(2) cells were cultured in DMEM F-12 media with 10% FBS and 1% pen-strep, COLO858 cells were cultured in RPMI with 5% FBS, 1% pen-strep and 1% sodium pyruvate, and K562 cells were cultured in RPMI with 20% FBS and 1% pen-strep. Cells were treated with 1 mM DFMO, 1 mM putrescine, 0.5 µM GC7, 1 µM UNC0368, 0.5 mM octyl-R-2HG, 25 µM JG2016, 0.5 µM Vorinostat, or 1 µM EPZ5676. For differentiation experiments, confluent 10T cells were exposed to chondrocyte differentiation media (DMEM containing 1% FBS, 1% pen-strep, 3e-8M sodium selenite, 10 µg/mL insulin, 1e-8M dexamethasone, 10 µg/mL transferrin, and 100ng/mL BMP-2). Confluent C2C12 cells were exposed to myotube differentiation media (DMEM containing 1% pen-strep and 2% horse serum). K562 cells were exposed to erythrocyte differentiation media (RPMI containing 40µM hemin) Differentiation media was refreshed every two to three days. For growth curves, 15,000 cells were plated in triplicate, and counted using a Beckman-Coulter Multisizer 3. For viability analysis, cells were stained with Trypan Blue, and counted using a Countess 3 Automated Cell Counter (ThermoFisher). All lines were determined to be mycoplasma free using MycoAlert Detection Kit (Lonza).

### Western Blots

Protein lysates were harvested from cells on ice with 1X RIPA buffer supplemented with protease and phosphatase inhibitors (Thermo Fisher # 1860932 and Thermo Fisher # 78428, respectively). For experiments in which histone marks were assayed, lysates were harvested in 1X RIPA buffer with 10% SDS and sonicated with a Bioruptor Plus (Diagenode). Protein was separated on NuPAGE 4-12% Bis-Tris gels (Invitrogen NP0336), blocked in 5% non-fat milk, and incubated with primary antibodies. After washing and secondary antibody incubation, blots were incubated in 1-shot ECL solution (Kindle Biosciences LLC) and imaged with a ChemiDoc MP Imaging System (BioRad).

### EdU and OPP incorporation and flow cytometry

Cells were treated for 25 minutes with 20 µM EdU or 20 µM OPP (Thermo Fisher C10456). Following this pulse, EdU-labeled cells were harvested, washed three times with 1% BSA in PBS, and fixed with 4% paraformaldehyde (PFA). After fixation, cells were permeabilized with 0.5% Triton X-100 in PBS, washed three times, and incubated in click chemistry reaction mix (2 mM CuSO4, 8 µM sulfo-azide dye, 20 mg/mL ascorbic acid in PBS) for 30 minutes in the dark. Cells were then washed, incubated in Hoechst (4 µg/mL), and EdU incorporation was assayed by flow cytometry (NovoCyte Quanteon).

OPP-pulsed cells were fixed in ice-cold methanol for 10 minutes at –20°C, washed in 1% BSA in PBS, and permeabilized with 0.5% Triton X-100. Cells were stained with homemade click chemistry reaction mix for thirty minutes, washed, and analyzed by flow cytometry (NovoCyte Quanteon).

### Gas chromatography mass spectrometry

GC-MS data collected over a time course of zebrafish embryonic development was analyzed from Dhillon et al., 2019^61^.

For extraction, chromatography and mass spectrometry analysis, cells were harvested in 80% Methanol, Optima LC/MS Grade, (Fisher Chemical) overnight at -80°C and dried using a Speed Vac Vacuum Concentrator (Savant) attached to a Universal Vacuum System (Savant). Samples were resuspended in an extraction buffer (10% NaCl solution made pH 1 with 0.5 M HCl) and polyamines were extracted with diethyl ether. Polyamines were derivatized by extracting two times with 15% ethyl chloroformate in diethyl ether and dried under a nitrogen stream. After another round of derivatization with trifluoroacetic anhydride in ethyl acetate, samples were dried under a nitrogen stream and resuspended in ethyl acetate. Polyamines were measured with the Agilent 8890/5977C gas chromatography/mass selective detector (GC/MSD). For LCMS, dried metabolites were resuspended in water with N-octyl-amine internal standard (0.025 µM) with 5 µL of 1M sodium bicarbonate (pH 9). Samples were derivatized with isobutyl chloroformate for 15 minutes at 37°C followed by diethyl ether extraction and drying of the organic phase under nitrogen stream. Samples were resuspended in 1:1 acetonitrile:water and 1 µL was analyzed on a Vanquish Duo UHPLC coupled to an Orbitrap ID-X Tribrid mass spectrometer (Thermo Fisher Scientific) via a heated electrospray ionization (HESI) source. Samples were injected in a randomized order to minimize batch effect. Samples were resolved by Reverse Phase Chromatography using an Acquity UPLC BEH C18 column (2.1 x 150 mm, 1.7 µM, Waters) maintained at 40 °C. The mobile phases consisted of (A) 0.1% formic acid in water, and (B) 0.1% formic acid in acetonitrile. Mobile phases were delivered as follow: from 0 to 3 minutes, 0% B (300 µL/min); from 3 to 16 minutes, 0% to 40% B linear gradient (300 µL/min); from 16 to 19 minutes, 40% to 100% B linear gradient (300 µL/min); from 19 to 21 minutes, 100% B (300 µL/min); from 21 to 21.9 minutes, 100% to 0% B linear gradient (300 µL/min); from 21.9 to 22 minutes, 0% B (300 µL/min); from 22 to 23.6 minutes, 0% B (500 µL/min); from 23.6 to 24 minutes, 0% B (300 µL/min). Eluates were ionized using a HESI source alternating between +3.5 kV to -2.5 kV potential (capillary temperature= 250 °C, sheath gas flow rate= 35 a.u., auxiliary gas flow= 5 a.u., and sweep gas flow= 1 a.u.). Ions were measured over a m/z range of 100-1000 using the orbitrap detector at 60,000 resolution (profile mode). The automatic gain control (AGC) target was set to standard with automatic maximum injection time control. Spectral data were processed using FreeStyle software (Thermo Fisher Scientific) and Skyline v.24.1 (McCoss Lab, University of Washington; MacLean B. et al., 2010, PMID: 20147306). For LC-MS experiments, quantitative analysis was performed on raw peak areas of standard-validated metabolites meeting the following criteria: coefficient of variation (CV) less than 30% in pooled quality control samples, signal-to-noise ratio (S/N) greater than 3, mass accuracy within 5 ppm, and absence of detectable signal in blank controls.

### RNAseq, scRNAseq analysis, ATACseq, and MNase digestion

RNAseq data was analyzed from the following repositories: GSE129957, GSE99399, GSE106292, GSE122380, GSE42519, and the LIBD Stem Cell Browser^62^.

RNA was harvested from cells (RNeasy Mini Kit, Qiagen 74104), and quality control was performed by Tapestation analysis (Agilent). Libraries were prepared with the TruSeq Stranded mRNA Library Prep Kit (Illumina) and sequenced on an AVITI system (Element Biosciences) using paired-end sequencing. bases2fastq was used to demultiplex the data, and reads were mapped to the mouse genome (GRCm10) using kallisto. Count matrices were analyzed in R using DESeq2.

For single cell analysis of erythroid differentiation, the count matrix was downloaded from GSE196347 containing transcriptomes from sorted CD71+ murine bone marrow sequenced on an Illumina NovaSeq and quantified using the Cell Ranger workflow. Dimensionality reduction and visualization were conducted by first learning the latent representation of the data using scVI^63^, a tool that models gene expression using a deep neural network. The model was run with 1 hidden layer, a latent space of dimension 10, and a dropout rate of 0.1. The scVI latent variables were used as input to the Scanpy UMAP function (sc.tl.umap) to produce a low-dimensional visualization of the data. Leiden clustering (sc.tl.leiden) was performed using a resolution of 0.5 to capture broad cell types, generating nine clusters. Each cluster was interrogated for expression of Hba-a1, Gypa, Slc4a1, and Tfrc. Two clusters were removed due to high mitochondrial gene expression or lack of Hba-a1 expression. Modeling of the data was repeated after cluster removal.

To reconstruct the progression of cells through erythropoiesis, PAGA^64^ was used to infer a trajectory (sc.pl.paga). A ‘root cell’ was determined based on the expression patterns of Tfrc, Klf1, Gata1, Xpo7, Tmcc2, and Bpgm, indicating the cluster at the start site of the trajectory^65–69^. The diffusion pseudotime was calculated using sc.tl.diffmap and sc.tl.dpt. Gene expression plotted against pseudotime was smoothed using the LOWESS (Locally Weighted Scatterplot Smoothing) tool in the statsmodels module. Confidence intervals were calculated by determining the standard deviation of 50 curves generated from a sampling of 500 data points (with replacement). The track plot was generated using sc.pl.tracksplot, with the groups (Ery1, Ery2, Ery3, Ery4, and Ery5) reflecting the Leiden clusters in order of pseudotime. Expression of essential erythropoietic genes and genes specific for hematopoietic progenitors, myeloid cells, lymphoid cells, and megakaryocytes were used to validate the generated pseudotime trajectory.

ATACseq library preparation was carried out as previously described^70^ and included a spike-in of 0.1 ng E. coli DNA per sample (EpiCypher # 18-1401). Libraries were amplified with NEBNext Multiplex Oligos (Illumina), and quality control was performed with Tapestation (Agilent) and KAPA library quantification (Roche). Libraries were sequenced on an AVITI system (Element Biosciences) using paired-end sequencing, and quality control was performed using FASTQC. Reads were trimmed using Trimgalore, and aligned to GRCm10 and the NCBI E. coli K12 genome using Bowtie2. Sorting and duplicate removal was performed using Samtools and Picard, and peaks were called with MACS2. Depth normalization to E. coli internal standards, and differential peak analysis, was performed using DiffBind. Peaks were annotated with ChipSeeker and CHIPpeakAnno, and reads were indexed for visualization with Samtools.

For MNase digestion experiments, nuclei from 147,000 cells per sample were incubated in 1 U MNase (ThermoFisher, EN0181) per million cells for 15 minutes at 37°C. DNA was extracted and quantified on a Qubit (ThermoFisher). 3 ng DNA per sample was analyzed with a high sensitivity Tapestation (Agilent).

### Polyamine probe and microscopy

Putrescine, spermidine, and spermine azide probes were synthesized as previously described^71^, without the addition of the BoDiPY moiety. Probes were resuspended in media at 10 µM and applied to live cells for two hours. Cells were then washed, fixed with 4% paraformaldehyde, permeabilized with 0.1% Triton-X in PBS, and incubated in homemade click chemistry reaction cocktail (2 mM CuSO4, 50mM THPTA, 8 µM alkyne-conjugated fluorophore, 20 mg/mL ascorbic acid) for 30 minutes. Following click chemistry reactions, cells were stained with Hoechst or DAPI, and imaged on a Zeiss LSM 780 confocal microscope.

For zebrafish probe experiments, AB wildtype embryos were manually dechorionated at the four-cell stage and moved into water containing 10 µM polyamine probe for 2 hours. Embryos were then fixed for 20 minutes with 4% PFA, permeabilized with 0.1% Triton X-100 in PBS, incubated in homemade click chemistry reaction cocktail with an alkyne-conjugated fluorophore for 30 minutes, and then DNA was stained with Hoechst. Yolk was removed by manual dissection and embryos were mounted for imaging on a Zeiss LSM 780 confocal microscope. The Institutional Animal Care and Use Committees of Columbia University approved all zebrafish experiments.

### In vitro enzymatic assays

All *in vitro* reactions were carried out with 1 µg nucleosome or histone protein substrate and 2.8 µg enzyme in 40 µL reaction buffer. Reactions were halted by addition of NuPAGE sample buffer, boiled at 95°C for 5 minutes, and then split in two and run on separate NuPAGE 4-12% Bis-Tris gels for analysis by western blot. *In vitro* demethylase reaction buffer contained 50 mM Tris-HCl pH 8.0, 5% glycerol, 2 mM ascorbate, 1 mM alpha-ketoglutarate, 100 µM FeSO4, 50 mM KCl, and protease inhibitor. *In vitro* demethylase reactions were carried out at 37°C for two hours. *In vitro* methyl-transferase reaction buffer contained 50 mM Tris-HCl pH 8.0, 5 mM MgCl2, 10 µM SAM, 4 mM DTT, and protease inhibitor. *In vitro* methyl-transferase reactions were carried out at 37°C for one hour. *In vitro* deacetylase reaction buffer contained 25 mM Tris-HCl pH 8.0, 50 mM NaCl, 1 mM DTT, 5 mM NAD+ and protease inhibitor. *In vitro* deacetylase reactions were carried out at 37°C for 15 minutes. *In vitro* acetylase reaction buffer contained 50 mM Tris-HCl pH 8.0, 5% glycerol, 1 mM DTT, 50 mM NaCl, 100 µM acetyl CoA, and protease inhibitor. *In vitro* acetylase reactions were carried out at 37°C for one hour.

Trypsin tail digestion reactions were carried out with 2 µg nucleosome or histone protein substrate and 10 ng trypsin in 10 µL reaction buffer. Reactions were halted by addition of Tricine sample buffer, boiled at 95°C for 10 minutes, and run on a Novex Tricine 16% mini protein gel. Gels were stained with Imperial protein stain. Trypsin digestion buffer contained 20mM MOPS, 1mM EDTA, and 1mM DTT.

To assess nucleosome integrity after polyamine incubation, mononucleosomes were incubated for one hour at 37°C in the presence of 100 µM putrescine, spermidine, or spermine. A digested control sample was processed with the Zymo Research Clean and Concentrate Kit. Samples were run on a 1.5% agarose gel, DNA was stained with ethidium bromide, and the gel was imaged on a ChemiDoc MP Imaging System (BioRad).

### Histone purification and nucleosome reconstitution for NMR

Histones and DNA were purified and reconstituted as outlined in Dyer et al^72^. Unmodified human histones (H2A.1 Uniprot accession P0C0S8, H2B.1C Uniprot accession P62807, H4 Uniprot accession P62805, and H3.2 Uniprot accession Q71DI3 with C110A and G102A) were expressed in Rosetta 2 (DE3) pLysS (Novagen) or BL21 (DE3) (New England Biolabs) chemically competent E. coli. Unlabeled growths were carried out in LB and induced at OD600∼0.4 with 0.2 mM (for H4) or 0.4mM (for H2A, H2B, and H3) IPTG for 3-4 hours. ^15^N-isotopically enriched H3 was grown in M9 minimal media supplemented with vitamin (Centrum) and 1 g L-^15^NH Cl. Histones were extracted from inclusion bodies and purified via ion exchange chromatography.

Histones were validated using ESI mass spectrometry (Fig. S6C). For DNA purification, a plasmid containing 37 repeats of the 147 bp Widom 601 sequence (ATCGAGAATC CCGGTGCCGA GGCCGCTCAA TTGGTCGTAG ACAGCTCTAG CACCGCTTAA ACGCACGTAC GCGCTGTCCC CCGCGTTTTA ACCGCCAAGG GGATTACTCC CTAGTCTCCA GGCACGTGTC AGATATATAC ATCCGAT) was amplified in E. coli and purified via alkaline lysis methods. The 601 repeats were released by cleavage with EcoRV and purified from parent plasmid by polyethylene glycol precipitation. Nucleosome reconstitutions were prepared by refolding tetramer (with equimolar ratios of H3 and H4) and dimer (with equimolar ratios of H2A and H2B) separately. Tetramer, dimer, and 601 DNA were then mixed together at a 1:2.2:1 molar ratio. Both mixtures were then desalted using a linear gradient from 2 M to 150 mM NaCl over 36-48 hours, followed by dialysis against 0.5xTE. Samples were then purified with a 10-40% sucrose gradient, which separates residual free 601 DNA and any subnucleosomal particles (i.e. tetrasome and hexasome). Native- and SDS-PAGE were used to assess the formation of nucleosomes along with histone composition (see Fig. S6A). Bands were visualized with ethidium bromide or Coomassie for native and denaturing gels, respectively. Nucleosomes were then dialyzed into 20 mM MOPS pH 7, 100 mM NaCl, 1 mM DTT, 1 mM EDTA for the purposes of NMR titrations. Concentrations were determined via UV-vis spectroscopy using the absorbance from the 601 DNA (calculated ε260 = 2,312,300.9 M-1cm-1). Samples were diluted into 2 M NaCl prior to concentration measurements in order to promote nucleosome disassembly for more accurate concentration determination.

### NMR Titration

^1^H,^15^N heteronuclear single quantum coherence (^1^H^15^N-HSQC) spectra of ^15^N-labelled ^15^N-H3-NCP (128 µM) in 20 mM MOPS pH 7, 100 mM NaCl, 1 mM DTT, 1 mM EDTA, and 10% D_2_O were measured at varying spermidine trichloride concentrations. All the HSQC spectra were measured at 37°C on a Bruker Avance Neo 600 MHz spectrometer equipped with a room temperature probe. The NMR spectra were processed in NMRPipe and analyzed using Sparky^73^. The normalized chemical shift difference (Δδ) was calculated using the following equation:

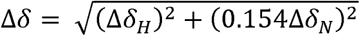

Where ΔδH and ΔδN are the observed changes in the ^1^H and ^15^N chemical shifts, respectively, upon addition of spermidine or spermine trichloride.

### NMR ^15^N Relaxation analysis

The backbone ^15^N longitudinal relaxation rates (R1), and transverse relaxation rate (R2) of ^15^N-H3 -NCP, in both the apo and spermidine trichloride or spermine bound-form were measured on a Bruker NEO 600 MHz spectrometer at 371, using gradient enhanced phase-sensitive HSQC pulse sequences (hsqct1etf3gpsitc3d and hsqct2etf3gpsitc3d). The ^15^N-H3-NCP (128 µM) sample and the ^15^N-H3-NCP sample with 512 µM spermidine or spermine were prepared in a buffer containing 20 mM MOPS (pH 7), 100 mM NaCl, 1 mM DTT, 1 mM EDTA, and 10% D O. The ^15^N-R1 experiments were conducted with relaxation delays of 25, 200, 400, 800, 1250, and 1750 ms. In the ^15^N-R2 experiments the relaxation delays were set to 15.68, 31.36, 47.04, 62.72, 94.08, and 109.76 ms. The ^15^N-R1, and ^15^N-R2, experiments were processed in NMRPipe. The R1 and R2 relaxation rates were determined by least-square fitting of the amide peak intensities to a single exponential decay function. Subsequently, the rotational correlation time, *τ*_c_ was calculated from R1 and R2 using the following equation:

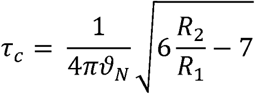

Where *ϑ*_N_ is the resonance frequency of ^15^N.

The error in R1 and R2 were propagated to R2/R1, and then to *τ*_c_ using standard error propagation.

### Reprogramming

Mouse embryonic fibroblasts (MEFs) expressing either doxycycline-inducible OSK or OSKM were reprogrammed as previously described^74^. Briefly, cells were seeded on a layer of feeder MEFs, then transferred into induction media (DMEM with 15% FBS, 1% non-essential amino acids, 1% GlutaMAX, beta-mercapto-ethanol and 1000 U/mL LIF) containing 1 mg/mL doxycycline and either 1 mM DFMO,1 mM putrescine, or 1 µM A485 (HATi). Media was refreshed every two days for twelve days, and then cells were moved into induction media without doxycycline for four days. Doxycycline-independent GFP-positive colonies were counted manually, and then live cells were dissociated for analysis by flow cytometry on a NovoCyte Quanteon.

## Acknowledgements

We thank C. Chio (Columbia University) and C. Lu (Columbia University) for antibody reagents and E. Mikhail Yohannan Cheria (Columbia University) for assistance with GCMS. Three and four factor reprogrammable fibroblasts were a gift from M. Stadtfeld (Weill Cornell Medical College). Flow cytometry and confocal microscopy were performed in the Columbia Stem Cell Initiative Flow Cytometry Institutional Core facility and the Columbia Medicine Microscopy Core. The histone acetyl-transferase inhibitor JG2016 was a gift from J. Gruber (UT Southwestern Medical Center). We thank C. Chio, C. Lu, S. Sternberg and L. Johnston for critical reading of the manuscript.

## Author Contributions

Conceptualization: MEB, JMS

Methodology: MEB, MRR, GF, AJS, SO, AG, RS, JC, SV, MHK, JS, SA, HAF, CM

Investigation: MEB, MRR, GF, AJS, SO, AG, RS, JC, MHK, SA, HAF, VSS, EMS

Data Curation: MEB, MRR, GF, AJS, SV, MHK, JS, SA, VSS, EMS, CM, JMS

Formal Analysis: MEB, MRR, GF, AJS, SV, JS, ADV, CM, JMS

Visualization: MEB, CM, JMS

Funding Acquisition: MEB, CM, JMS

Project Administration: JMS

Supervision: JMS

Writing – original draft: MEB, JMS

Writing – review & editing: MEB, JMS

## Data and Materials Availability

All data and materials are available upon request. Sequencing data are deposited with the NCBI GEO (accession pending).

