## Supplemental Figures and Figure Legends for "Polyamines regulate cell fate by altering the accessibility of histone tails"

#### Supplementary Figure Legends

##### Supplemental Figure 1

A-B) ODC1 expression from publicly-available RNAseq data collected over time courses of granulopoiesis and *in vitro* cardiomyocyte differentiation. For cardiomyocyte differentiation all comparisons to day 1 $p \leq 0.0001$.

C-H) Expression of polyamine biosynthesis genes across time courses of osteoblast, chondrocyte, myotube, adipocyte, erythrocyte and granulocyte differentiation. Abbreviations for granulocyte differentiation time course: HSC – hematopoietic stem cell; MPP – multipotential progenitor; CMP – common myeloid progenitor; GMP – granulocyte monocyte progenitor; early PM – early promyelocyte; late PM – late promyelocyte; MY – myelocyte; MM – metamyelocyte; BC – band cell; PMN – polymorphonuclear cell.

I-J) Quantification of putrescine and spermidine abundance by gas-chromatography mass-spectrometry over the first 96 hours of zebrafish development. P values calculated by Tukey's test and shown only for decrease from maximum at 48 hours.

##### Supplemental Figure 2

A) Western blot for ODC1 in Rosa26 and ODC1 gRNA transduced 10T1/2 single cell clones.

B) Quantification of spermidine and spermine by GC-MS in control (sgRosa) and ODC1 knockout (sgODC1) cells following three days of treatment in putrescine-free media. Polyamine content was normalized to cell number. P values calculated by one-tailed t-test.

C) Quantification of putrescine, spermidine and spermine by GC-MS in control (sgRosa) cells following three days of treatment in media containing vehicle, 1 mM DFMO, or 1 mM putrescine. Polyamine content was normalized to cell number. P values calculated by t-test with Bonferroni correction.

D) Cell viability after three days of treatment in vehicle, 1 mM DFMO, or 1 mM putrescine was assessed by Trypan Blue staining. Dashed line indicates 100%. P values calculated by t-test with Bonferroni correction.

E) Cell cycle analysis by EdU incorporation assay. Following three days of treatment in vehicle, 1 mM DFMO, or 1 mM putrescine, cells were pulsed with 20 μM EdU for 25 minutes, fixed, and EdU fluorescence was assessed by flow cytometry. Cell cycle was determined by comparing EdU incorporation to DNA content (by Hoechst staining). Data presented are the averages of three biological replicates. P values correspond to t-test with Bonferroni correction for the S phase population.

F) Gene set enrichment analysis showing top terms enriched in low-polyamine samples (control cells treated with DFMO, and ODC1 knockout cells treated with vehicle or DFMO) as compared to high-polyamine samples (control cells treated with vehicle or putrescine, and ODC1 knockout cells treated with putrescine).

G) Gene set enrichment analysis showing top terms enriched in high-polyamine samples (control cells treated with vehicle or putrescine, and ODC1 knockout cells treated with putrescine) as compared to low-polyamine samples (control cells treated with DFMO, and ODC1 knockout cells treated with vehicle or DFMO).

H) Western blot of adipocyte differentiation marker PPAR-gamma in vehicle, DFMO and putrescine-treated c10T1/2 cells in growth or adipocyte differentiation conditions.

**Supplemental Figure 3**

A) SK-N-BE(2) neuroblastoma and K562 leukemia cells were plated in the indicated treatments (1 mM DFMO, 1 mM putrescine) for either four or five days, respectively, and cell number was quantified. Data presented are mean +/- standard error of three biological replicates. P values are shown on plot; Dunn's test.

B) Cells were pulsed with O-propargyl-puromycin (OPP) for 25 minutes following three days of the indicated treatments (1 mM DFMO, 1 mM putrescine). Positive control cells were treated with 10 mg/mL cycloheximide for 25 minutes. OPP incorporation was quantified by flow cytometry.

C) Frequency of cells with incorporated OPP per condition. Data presented are mean +/- standard error of three biological replicates. No significant differences by ANOVA (sgRosa $p = 0.184$; sgODC1 $p = 0.093$).

D) Cells were treated with 1 mM DFMO, 1 mM putrescine, or 50-500 μM thapsigargin, and eIF2a phosphorylation was assessed by western blot.

E) COLO858 melanoma and SK-N-BE(2) cells were treated with 0.5 μM GC7 for three days, then hypusine levels and differentiation marker expression were determined by western blot.

**Supplemental Figure 4**

A) Confocal images of cells treated with no probe, control probes (D-alanine and L-Ornithine) or putrescine probe and subjected to click-chemistry mediated addition of a fluorophore. Scale bar 50 μM.

B) Immunofluorescence confocal images of putrescine probe in control (sgRosa) or ODC1 knockout (sgODC1) cells following three days of treatment with putrescine-free media. Scale bars are 5 μM.

C) Structures of the spermidine and spermine azide probes.

D) Immunofluorescence confocal images of putrescine probe in U-2 OS osteosarcoma cells following three days of treatment with 1 mM DFMO or 1 mM putrescine. Scale bars are 5 μM.

E) Immunofluorescence confocal images of putrescine probe in RPE-1 retinal pigment epithelial cells following three days of treatment with 1 mM DFMO or 1 mM putrescine. Scale bars are 5 μM.

F) Top) Cells were treated with RNAse for thirty minutes, then RNA was extracted and quantified. P value corresponding to a t-test is shown. Bottom) Immunofluorescence confocal images of putrescine probe-treated 10T1/2 cells treated with/without RNAse prior to click-chemistry reaction.

G) Localization of putrescine probe in zebrafish embryos dechorionated for two hours after fertilization and exposed to the putrescine probe for two hours. Inset shows higher magnification of nuclei. Scale bars are 25 μM and 5 μM (insets).

**Supplemental Figure 5**

A) Control (sgRosa) and ODC1 knockout (sgODC1) cells were exposed to MNase digestion for 15 minutes. DNA was extracted, and fragment size was quantified by Tapestation. Relative abundance of fragment sizes is plotted.

B) Heatmap showing similarity between ATACseq samples.

C) Venn diagram of consensus peak overlaps (present in two out of three replicates) between ATACseq sample conditions.

D) Gene set enrichment analysis of peaks unique to DFMO-treated samples in transcriptional start sites.

E) Motif enrichment analysis of peaks which were uniquely accessible following three days of treatment in 1 mM DFMO.

F) Western blot showing abundance of indicated histone modifications in control and sgODC1 10T1/2 cells after three days of culture in media without putrescine.

G) Western blot showing H3K18ac and H3K9me2 abundance following cell cycle arrest in G1 by palbociclib treatment.

H) Dot blot showing levels of DNA 5mC in 10T1/2 cells following three days of treatment in indicated conditions (1 mM DFMO, 1 mM putrescine, 0.5 mM octyl-R-2HG).

I) H3K9me2 and H3K9me3 levels were assessed in 10T1/2 and SK-N-BE(2) cells by western blot after 24 hours of vehicle or 1 mM DFMO treatment, followed by 24 hours of methyltransferase inhibitor (1 μM UNC0368) or demethylase inhibitor (0.5 mM octyl-R-2HG), in the presence or absence of DFMO.

J) Western blot showing *in vitro* acetyl-transferase activity of Gcn5 on nucleosome substrate in the presence of indicated charge-matched concentrations of putrescine, spermidine and spermine.

K) Western blot showing *in vitro* acetyl-transferase activity of Gcn5 on histone H3 substrate in the presence of indicated charge-matched concentrations of putrescine, spermidine and spermine.

L) Western blot showing *in vitro* H3K9 demethylation activity of Kdm4a on histone H3 substrate in the presence of indicated charge-matched concentrations of putrescine, spermidine and spermine.

M) Western blot showing *in vitro* Sirt6 deacetylase activity on H3K18 acetylated histones in the presence of charge-matched concentrations of putrescine, spermidine and spermine.

N) Ethidium bromide-stained agarose gel electrophoresis showing recombinant nucleosomes incubated in vehicle or indicated polyamine concentrations. Proteinase K digested control shows free DNA migrating at ~150 bp.

O) 10T1/2 cells were treated with vehicle, 1 mM DFMO or 1 mM putrescine for 24 hours, and then treated with 1 μM EPZ5676 to inhibit DOTL1 for 24 hours. H3K79me2 levels were measured by western blot.

P) Western blot showing H3K79me2 abundance following *in vitro* methyltransferase reactions. Recombinant DOT1L was incubated in the presence of unmodified mononucleosomes and 100 μM putrescine, spermidine, or spermine for one hour at 37°C. Reactions were quenched with SDS, boiled, and assessed by western blot.

**Supplementary Figure 6**

A) Imperial stain of SDS-PAGE gel indicating digestion of intact nucleosomes in the absence or presence of indicated polyamines. 3 replicates shown as groups of 4 lanes following no trypsin control.

B) Imperial stain of SDS-PAGE gel indicating digestion of intact nucleosomes in the absence or presence of indicated polyamines and polyamine mix.

C) Imperial stain of SDS-PAGE gel indicating digestion of H3 histone protein in the absence or presence of indicated polyamines.

D) Native (left) and SDS-PAGE (right) analysis of the $^{15}$N-H3-NCP confirming nucleosome reconstitution.

E) ESI mass spectrometry analysis of purified histones confirming the expected molecular mass (13775 Da for H2A, 13960 Da for H2B, 15454 Da for H3, and 11236 Da for H4) and that none are carbamylated due to urea purification.

F) Left) Full $^{1}$H,$^{15}$N-HSQC spectra of the $^{15}$N-H3-NCP in the absence (light blue) and presence (dark blue) of spermidine. Right) Full $^{1}$H,$^{15}$N-HSQC spectra of the $^{15}$N-H3-NCP in the absence (grey) and presence (green) of spermine.

**Supplementary Figure 7**

A) Oct4-GFP expression of OSKM murine embryonic fibroblasts reprogrammed as in Figure 6 with addition of HATi (histone acetyl-transferase inhibitor) or HATi plus putrescine quantified by flow cytometry.

B) Oil Red O absorbance in 10T1/2 cells subjected to an adipocyte differentiation protocol in indicated conditions. P-values correspond to Tukey test.

### Supplementary Figure 1

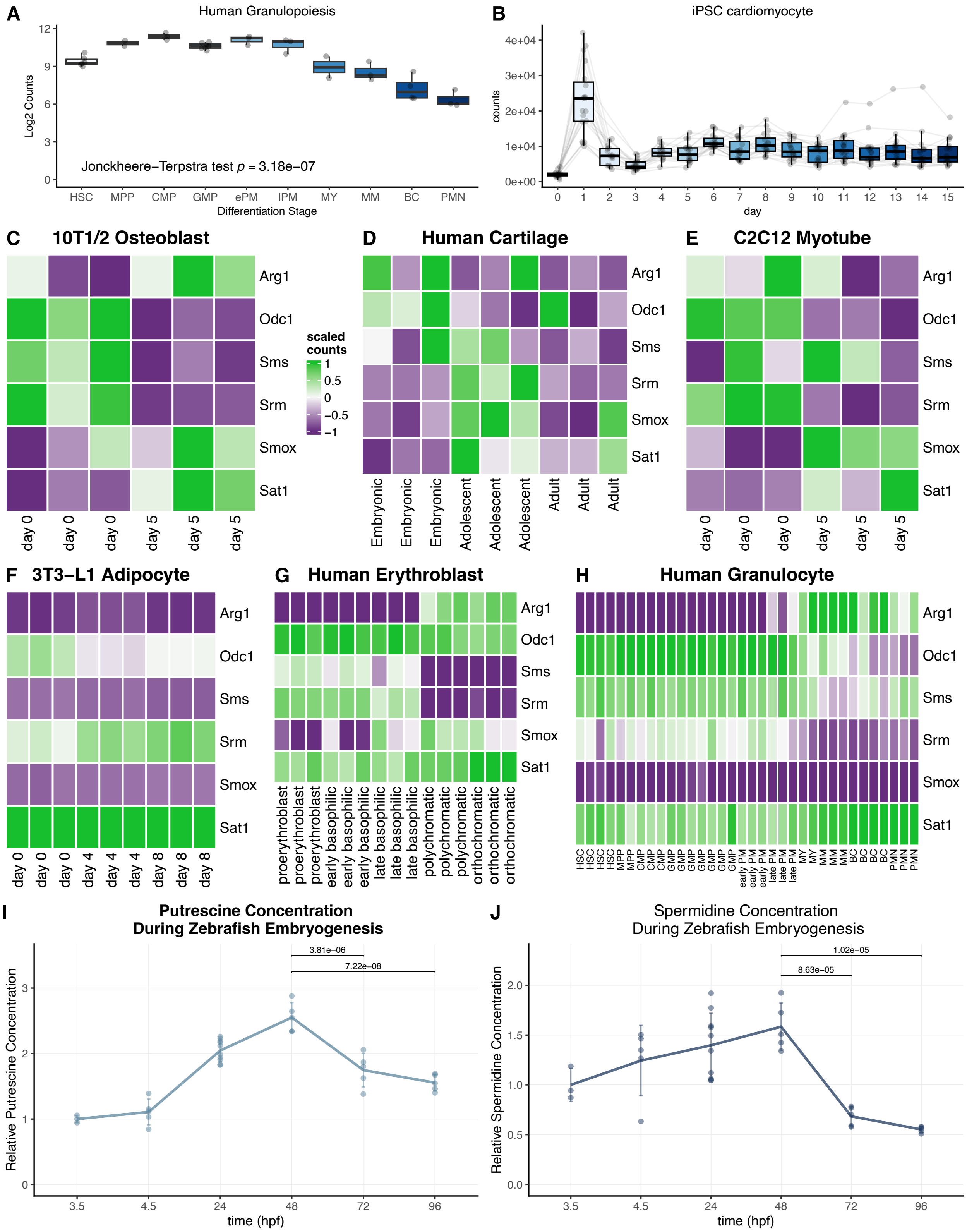

#### Supplementary Figure 2

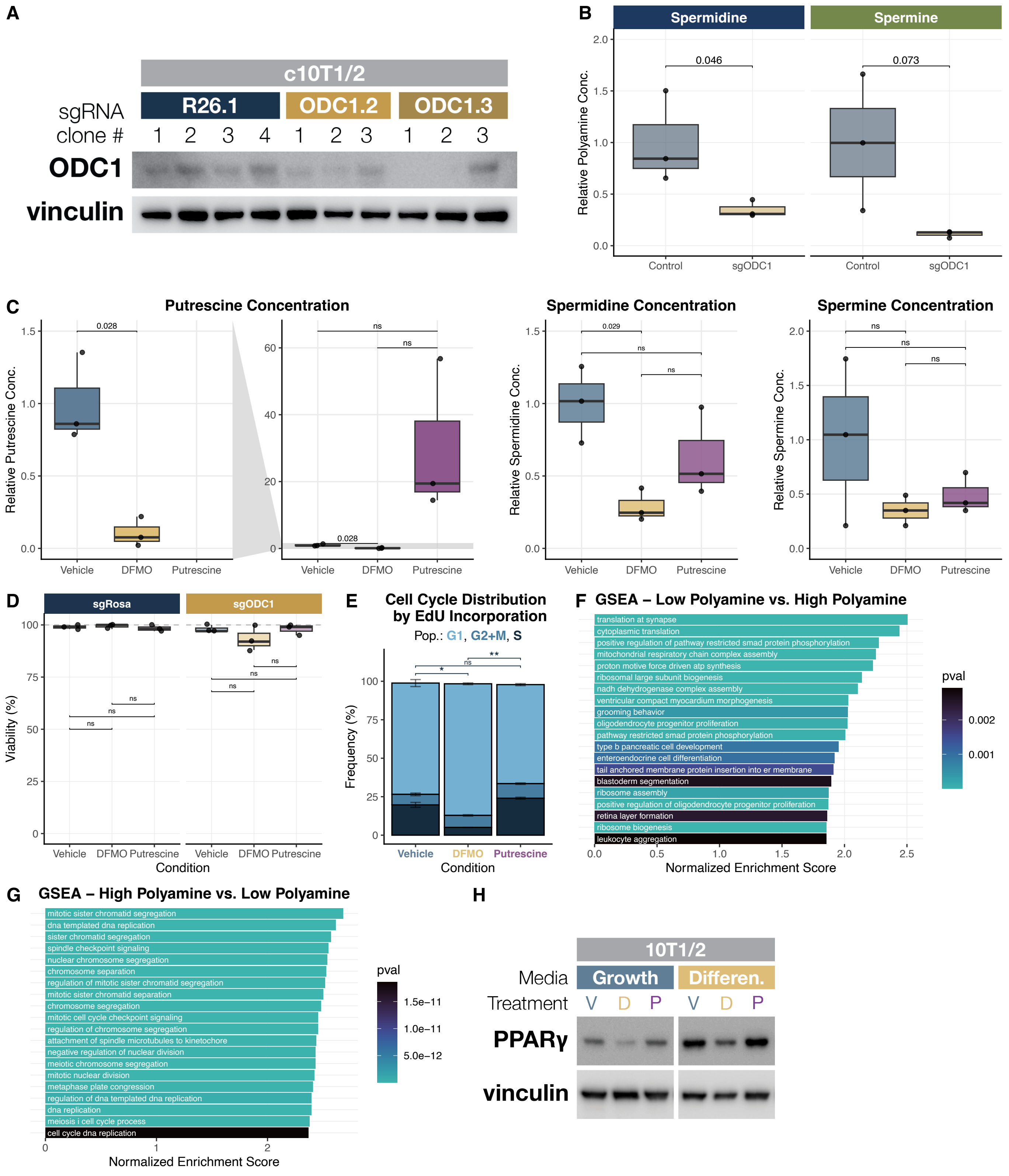

### Supplementary Figure 3

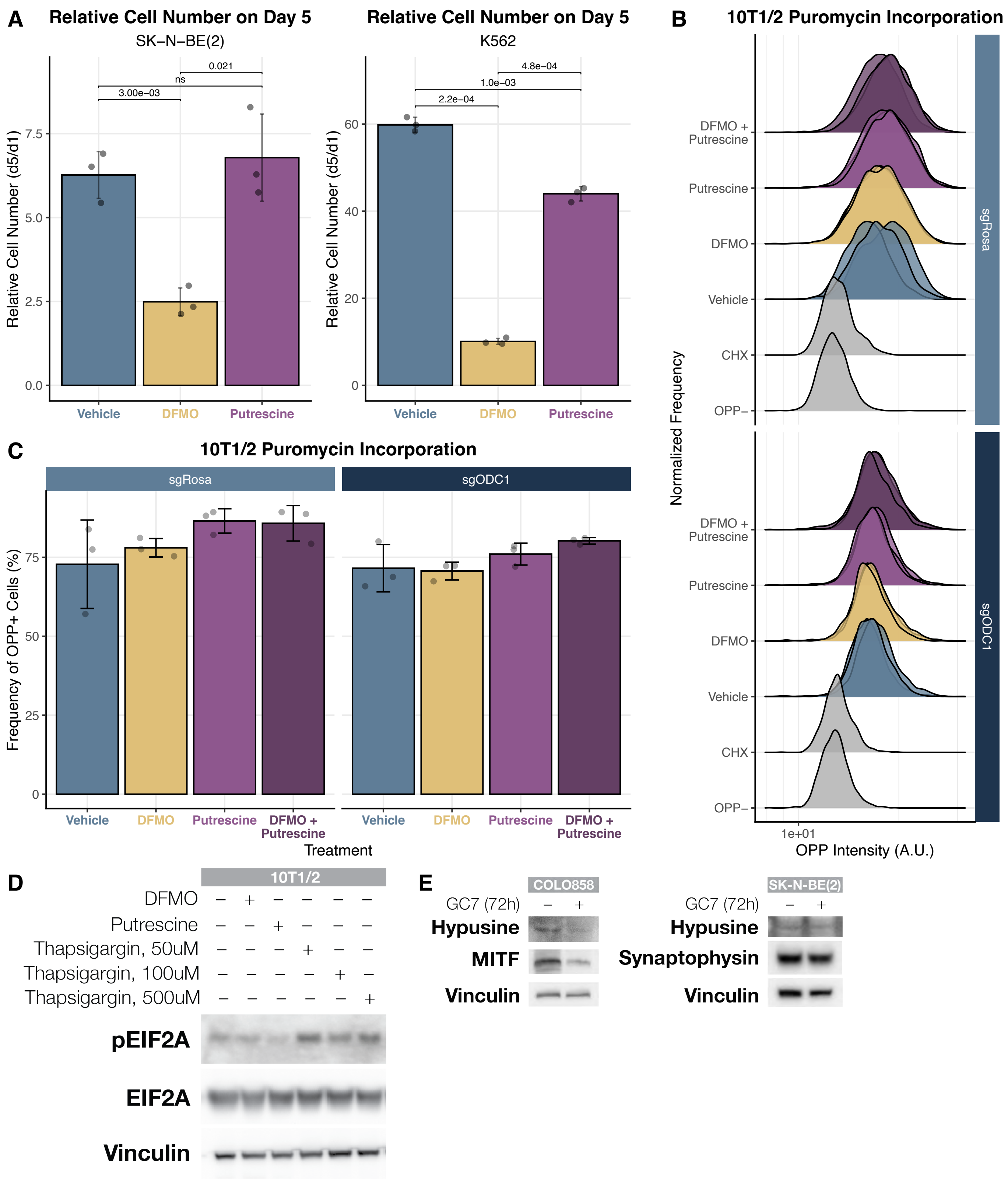

### Supplementary Figure 4

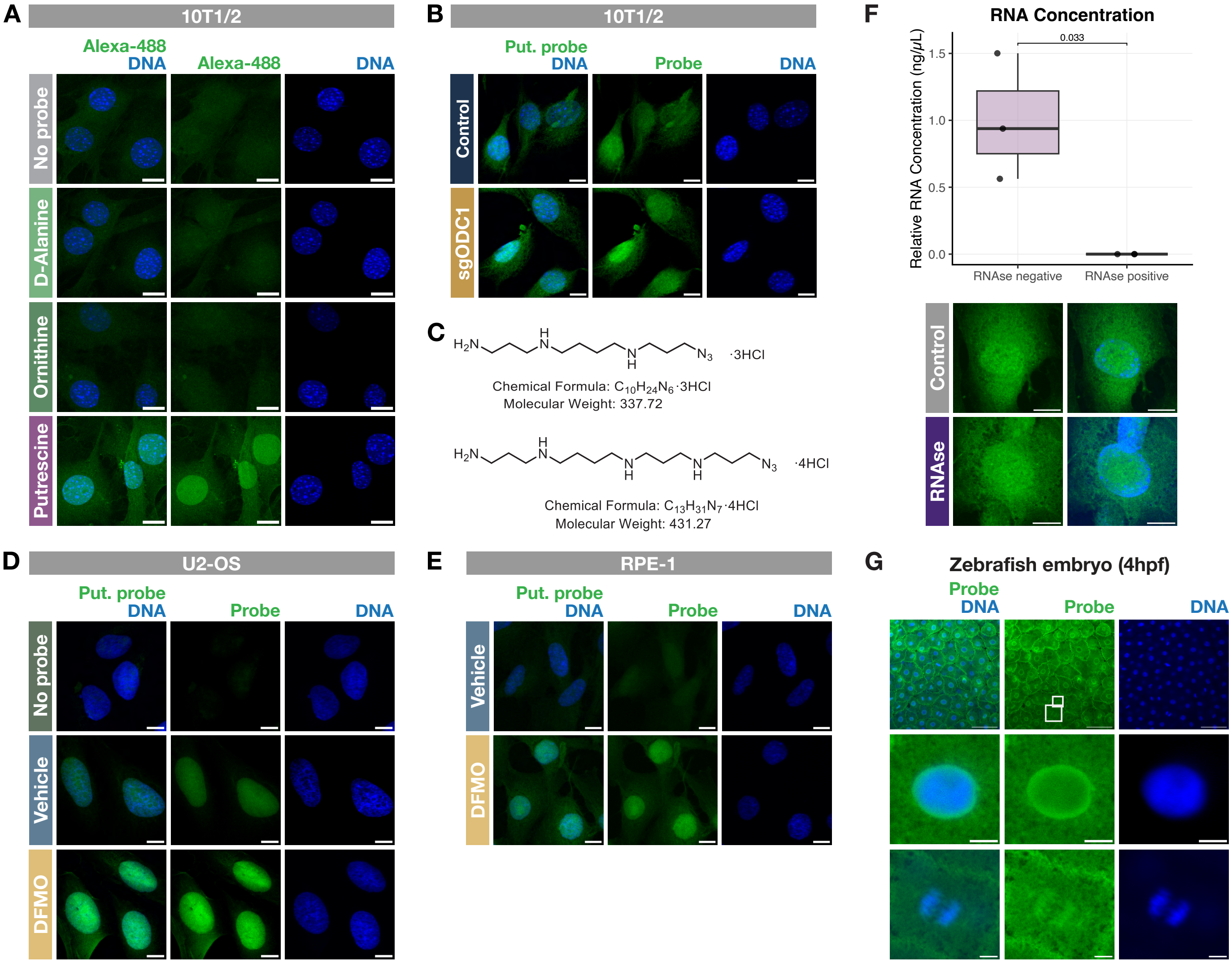

### Supplementary Figure 5

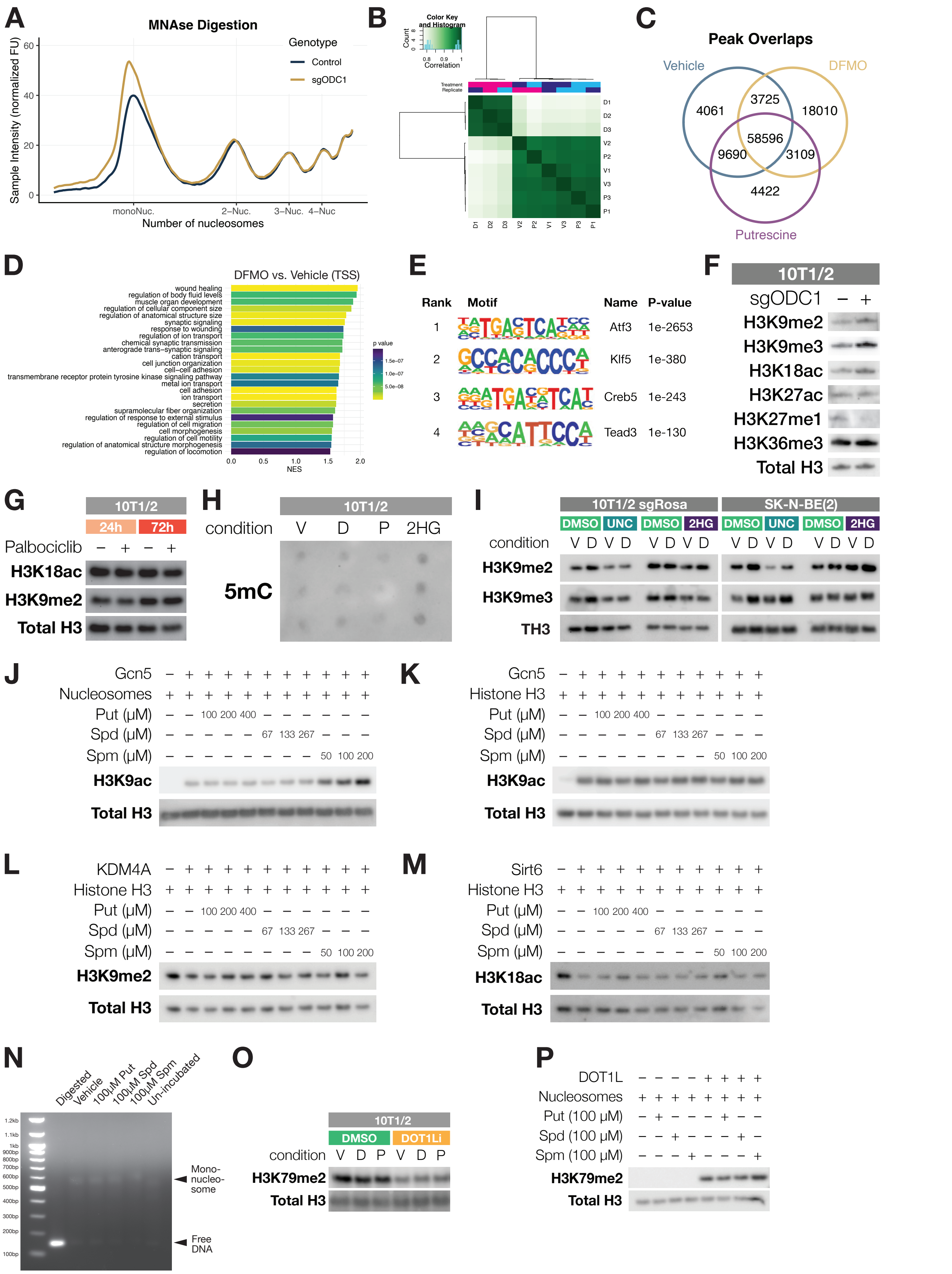

### Supplementary Figure 6

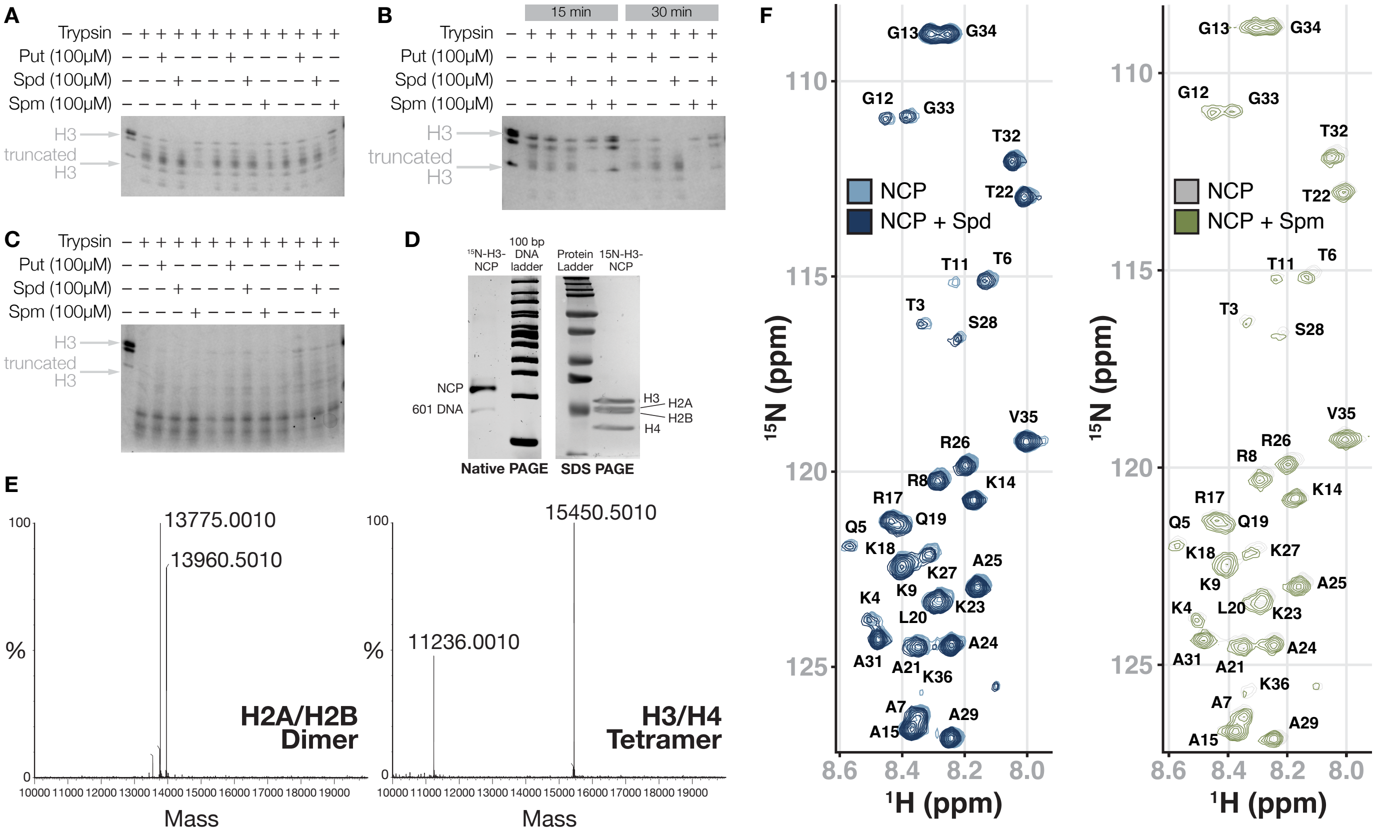

### Supplementary Figure 7

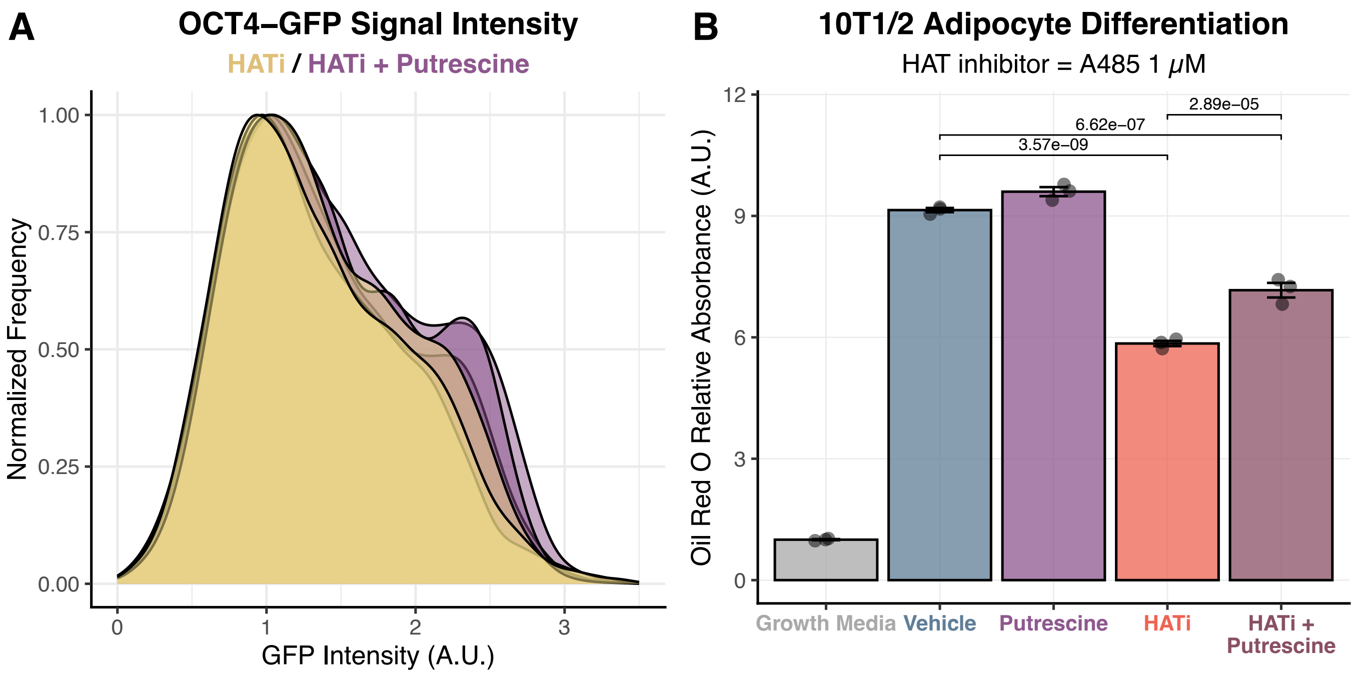
